# Fasting disrupts the InsP₆–HDAC3 axis to drive ER stress–mediated clearance of DNA-damaged cells and enforce tissue quality control

**DOI:** 10.64898/2026.05.24.727426

**Authors:** Sujan Chetterjee, Zachary Sin, Richard A. Van, Nguyen Tran, Tam Tran, Anuj Shukla, Lucile Marie-Paule Jeusset, George Koshkaryan, Kevin Saldana, Rayan Muneer, Loretta Viera Preval, Lingkun Gu, Faizan Bhat, Kevin Ritter, Katherine Huang, Kayci Huff-Hardy, Richard Rood, Anas Gremida, Martin Gregory, Chien Huan Chen, Mo Weng, Frank J Dekker, Parakkal Deepak, Henning J. Jessen, Kirk James McManus, Mira V. Han, Prasun Guha

## Abstract

Fasting drives metabolic adaptation but also elicits acute cellular stress. How this stress shapes tissue integrity is unknown. Here, we show that in the intestine, fasting depletes growth factor signaling, which triggers cellular stress. This response functions as a tissue quality-control checkpoint that selectively eliminates pre-existing DNA-damaged cells while sparing healthy counterparts. A short-term fast diminishes TGF-β signaling and elicits endoplasmic reticulum (ER) stress, driving DNA-damaged intestinal cells beyond an apoptotic threshold, thereby reducing the inflammatory burden. Mechanistically, loss of TGF-β signaling triggers FBXO22–Cullin1-mediated degradation of the inositol kinase IPMK, leading to depletion of inositol hexaphosphate (InsP₆). InsP₆ loss attenuates HDAC3 activity and initiates coordinated epigenetic and post-translational reprogramming, thereby increasing CDK5RAP3 abundance. Elevated CDK5RAP3 inhibits ribosomal RPL26 UFMylation, thereby amplifying ER stress and selectively licensing apoptosis in DNA-damaged cells. Collectively, fasting disrupts a TGF-β-InsP_6_-HDAC3 axis to drive ER stress-dependent clearance of DNA-damaged cells, enforcing tissue quality control.

## Introduction

Tissue integrity is maintained through surveillance mechanisms that identify and eliminate damaged cells before they compromise organ function. This requirement is particularly critical in rapidly renewing tissues such as the intestinal epithelium, where stem cell–derived progenitors proliferate continuously and differentiate along the crypt–villus axis^1^. The high proliferative activity of this tissue inherently increases the likelihood of spontaneous DNA lesions^2^, even in the absence of exogenous genotoxic stress. Although DNA damage can activate apoptotic pathways that eliminate compromised cells, the efficiency and magnitude of this response vary across physiological contexts and individuals, influenced by factors such as age, genetic background, and environmental exposures^3,4^. Consequently, cells harboring DNA damage may persist and accumulate within tissues, potentially increasing susceptibility to chronic inflammation and age-associated diseases^5^. These observations raise the question of how physiological signals may be engaged to selectively eliminate DNA-damaged cells and thereby preserve tissue integrity.

Fasting represents a metabolic state that profoundly reshapes cellular physiology^6^. While nutrient deprivation imposes systemic metabolic and cellular stress^7^, fasting followed by refeeding has been shown to enhance stem cell activity and promote tissue regeneration across multiple organs^8,9^. These observations raise the possibility that fasting may first alter epithelial cell composition before the regenerative phase begins. Because cells harboring DNA damage already experience intrinsic genotoxic stress, additional cellular stress imposed by fasting could selectively push these cells beyond an apoptotic threshold while leaving healthy cells largely unaffected. Such a mechanism would allow damaged cells to be removed prior to tissue renewal, thereby improving epithelial fitness. However, how fasting functions as a physiological signal that selectively eliminates DNA-damaged epithelial cells before the onset of regeneration remains unknown.

Here, we show that healthy human and mouse intestines harbor a substantial population of epithelial cells with DNA damage. Having observed that a brief fasting period eliminates these DNA-damaged cells, we sought to define the underlying mechanisms. We observed that fasting induces an endoplasmic reticulum (ER) stress–dependent program that selectively promotes apoptosis in DNA-damaged cells while sparing the healthy epithelium. Mechanistically, fasting disrupts a previously unrecognized growth factor-responsive pathway involving higher-order inositol phosphate (HOIP) metabolism and epigenetic modifications, thereby eliciting ER stress. Specifically, fasting depletes inositol hexaphosphate (InsP₆), attenuating histone deacetylase 3 (HDAC3) activity and reshaping both histone and non-histone acetylation to disrupt ribosome-associated proteostasis and induce ER stress. In cells harboring genomic damage, this stress response drives selective apoptosis. While our previous work established InsP₆ as an essential cofactor for HDAC3 activity^10^, the present study defines the physiological regulation of this axis and reveals its disruption during fasting.

Together, these findings establish fasting as a physiological mechanism of tissue quality control. By suppressing growth factor signaling, fasting disrupts the InsP₆-HDAC3 axis to reprogram chromatin and proteostasis, engaging ER stress across the epithelium. Although this stress is broadly induced, apoptotic execution is selectively restricted to DNA-damaged cells, thereby refining tissue composition without compromising epithelial integrity. This work not only reveals an unexpected physiological burden of DNA-damaged cells in healthy intestine, but also identifies a tractable mechanism by which transient nutrient deprivation can eliminate these compromised cells. More broadly, our study establishes a connection between growth factor levels, epigenetic regulation, and proteostasis, forming an axis that links these factors to genomic integrity. This finding suggests that fasting is not simply a metabolic adjustment but acts as a selective force that maintains the fitness of tissues.

## Results

### Short-term fasting selectively drives apoptotic clearance of pre-existing DNA-damaged intestinal epithelial cells

Highly proliferative tissues are intrinsically susceptible to spontaneous DNA double-strand breaks (DSBs)^2^, even in the absence of exogenous genotoxic stress. To determine whether this basal DNA damage is tissue-selective, we profiled DSB markers across major organs from healthy, untreated mice (10–16 weeks old). Immunoblot analysis of phospho-γ-H2AX (Ser139), a canonical DSB marker^11^, revealed prominent signal specifically in intestinal epithelial cells (IECs) from both ileum and colon, whereas liver, kidney, lung, pancreas, and brain (hippocampus and cortex) showed little to no signal **(Fig. 1A).** Phosphorylated Chk2 (pChk2), an independent marker of DSB signaling^12^, showed the same pattern, confirming selective enrichment of spontaneous DNA damage in IECs **(Fig. 1A).** To exclude basal inflammation as a confounding variable, we assessed cleaved IL-1β (mature) and found it to be undetectable in healthy untreated mice, in contrast to Dextran Sodium Sulfate (DSS)-treated animals used as a positive control for colonic epithelial injury **(Fig. 1B).** Thus, spontaneous DNA damage in IECs is not simply a consequence of overt inflammation.

**Figure 1.**
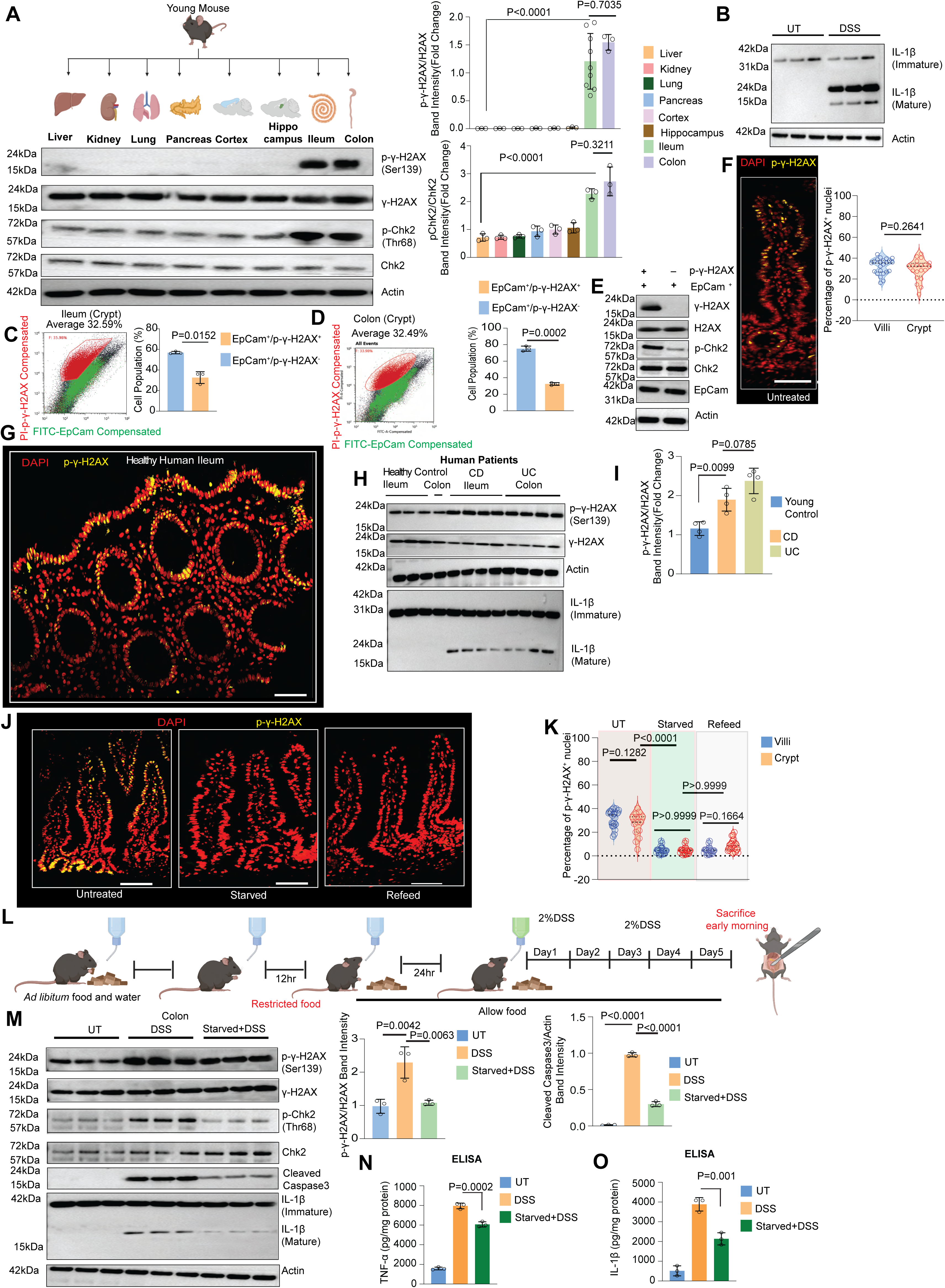
Short-term fasting promotes the apoptotic removal of DNA-dmaged intestinal epithelial cells, mitigating inflammation. **(A)** Immunoblot and densitometric analysis across indicated organs (n = 3-9; mean ± SD). Schematic created in BioRender (CHATTERJEE, S., 2026; https://BioRender.com/iz17aga). **(B)** Immunoblot shows mature and immature IL-1β levels in DSS-treated (2%, 5 days) and untreated mice colonic crypts (n ≥ 3). **(C–D)** Flow cytometry of ileal **(C)** and colonic **(D)** IECs shows EpCAM⁺/p-γ-H2AX⁺ and EpCAM⁺/p-γ-H2AX⁻ populations (n = 3; mean ± SD). **(E)** Immunoblot comparing FACS-sorted EpCAM⁺/p-γ-H2AX⁺ to EpCAM⁺/p-γ-H2AX⁻ cells (n = 3). **(F–G)** Immunohistochemistry detects p-γ-H2AX⁺ cells (yellow) in mouse ileum **(F)** and healthy human ileum tissue **(G)** (n = 4); merged signal shown with DAPI (pseudo-colored red); scale bar, 50 μm; quantification from 8 images per mouse (mean ± SD). Each dot is each snap, total 8 snapsX3 mice= total 24 dots. **(H–I)** Immunoblot **(H)** and densitometry **(I)** of human intestinal biopsies from healthy control ileum (n=3), colon (n=1), ulcerative colitis (UC), and Crohn’s disease (CD) (n = 4; mean ± SD). **(J–K)** Immunohistochemistry **(J)** and quantification of p-γ-H2AX⁺ cells **(K)** in untreated, starved, and refeed mouse ileum (n = 3; 300–600 nuclei per mouse; mean ± SD. Each dot is each snap, total 8 snaps X3 mice= total 24 dots). **(L)** DSS treatment scheme (BioRender; CHATTERJEE, S., 2026; https://BioRender.com/mtrovit). **(M)** Immunoblot and densitometry from untreated, DSS, and fasted+DSS mice colonic crypts (n = 3; mean ± SD). **(N–O)** ELISA shows TNF-α **(N)** and IL-1β **(O)** in indicated groups (n = 3; mean ± SD). Dots in all bar plots represent individual biological replicates.

We next quantified the abundance of DNA-damaged cells within the healthy intestinal epithelium by flow cytometry of purified EpCAM⁺ IECs stained for phospho-γ-H2AX (Ser139). In the ileum, approximately 32.6% of EpCAM⁺ cells were phospho-γ-H2AX^+^, indicating that nearly one-third of IECs harbor DNA damage at baseline **(Fig. 1C).** A similarly large phospho-γ-H2AX^+^ population (∼33.9%) was present in the colon **(Fig. 1D),** indicating that this feature is shared across intestinal compartments.

To validate these two populations biochemically, we flow-sorted EpCAM⁺/phospho-γ-H2AX⁺ and EpCAM⁺/phospho-γ-H2AX⁻ IECs and analyzed them by immunoblotting. Strong phospho-γ-H2AX and pChk2 signals were confined to the EpCAM⁺/phospho-γ-H2AX⁺ fraction, whereas the negative fraction lacked detectable DNA damage markers **(Fig. 1E).** Thus, the normal young-adult mouse intestine contains two major IEC populations distinguished by DNA damage status. All subsequent mouse experiments, unless otherwise specified, were performed in ileal IECs as a representative intestinal compartment.

We next examined the spatial distribution of DNA-damaged cells. Immunohistochemistry revealed phospho-γ-H2AX⁺ cells throughout the crypt-villus axis, without obvious compartmental restriction **(Fig. 1F).** DNA-damaged cells therefore reside in both proliferative and differentiated regions, indicating that the healthy intestinal epithelium contains a broad and widely distributed reservoir of damaged cells.

We then asked whether this phenomenon is conserved in humans. Analysis of ileal biopsies from healthy individuals (40–55 years old) obtained during routine colonoscopy revealed a robust phospho-γ-H2AX signal in normal epithelium, closely resembling the pattern seen in mice **(Fig. 1G, H).** Samples from Crohn’s disease (CD) and ulcerative colitis (UC) patients showed only moderately higher phospho-γ-H2AX levels than healthy controls **(Fig. 1H, I),** indicating that constitutive DSB accumulation is not restricted to disease. In parallel, cleaved IL-1β was undetectable in normal human epithelium but readily detected in CD and UC samples **(Fig. 1H),** demonstrating that healthy tissue is not inflamed despite carrying substantial DNA damage. These findings establish that spontaneous DNA damage is a physiological feature of the human intestinal epithelium.

Because DSBs can prime cells for apoptosis^13,14^, we next asked whether a physiological stress could accelerate clearance of DNA-damaged IECs. We first assessed basal apoptotic priming by measuring cleaved caspase-3 in sorted IEC populations. Cleaved caspase-3 was detectable at baseline in phospho-γ-H2AX⁺ cells but absent from the phospho-γ-H2AX⁻ fraction **(Supplementary Fig. 1A),** indicating that DNA-damaged IECs are pre-primed for apoptotic execution.

We then subjected mice to a 12h fast (water only), a physiological stress known to engage apoptotic pathways^15^. To separate nutrient stress from circadian effects, mice were fasted during their active phase (8:00 PM–8:00 AM). Strikingly, a single 12 h fast nearly eliminated phospho-γ-H2AX⁺ IECs, as assessed by flow cytometry **(Supplementary Fig. 1B),** immunoblotting **(Supplementary Fig. 1C),** and histology **(Fig. 1J,K).** Importantly, tissue architecture remained intact as assessed by H&E staining **(Supplementary Fig. 1D),** indicating selective removal of DNA-damaged cells rather than generalized epithelial injury. Notably, even after 24 h of refeeding following the fast, the regenerated epithelium remained largely devoid of DNA-damaged cells **(Fig. 1J, K).**

To determine whether this removal of DNA damaged cells reflects amplified apoptotic signaling in the pre-primed damaged population, we analyzed cleaved caspase-3 during the early phase of clearance. At 6 h of fasting, when phospho-γ-H2AX⁺ cell loss is initiated **(Supplementary Fig. 1E, F),** cleaved caspase-3 was markedly increased in the EpCAM⁺/phospho-γ-H2AX⁺ fraction relative to untreated controls **(Supplementary Fig. 1G)**. By contrast, the EpCAM⁺/phospho-γ-H2AX⁻ fraction showed only minimal caspase-3 cleavage even after fasting **(Supplementary Fig. 1G).** Thus, fasting selectively intensifies apoptotic signaling in DNA-damaged IECs while sparing undamaged cells.

Together, these data reveal that short-term (12 hours) fasting selectively drives the apoptotic clearance of preexisting DNA-damaged cells from the mouse intestine.

### Short-term fasting mitigates inflammation by reducing the burden of DNA-damaged cells

Pre-existing DNA damage can amplify inflammation through cGAS-STING and senescence-associated pathways^16^. Clearance of damaged cells before injury would therefore be expected to reduce inflammation. We next asked whether the removal of DNA-damaged IECs enhances epithelial resilience during epithelial injury. To test this, mice underwent a single 12 h fast followed by 24 h of refeeding before DSS (5 days) administration **(Fig. 1L).** This brief fasting-refeeding intervention conferred marked protection against DSS-induced acute colitis. Fasted-refed mice showed reduced DNA damage signaling, with lower phospho-γ-H2AX and pChk2 levels than continuously fed controls **(Fig. 1M).** DSS-induced epithelial apoptosis, measured by cleaved caspase-3, was prominent in continuously fed animals but markedly reduced in fasted-refed mice **(Fig. 1M).** In parallel, inflammatory responses were blunted, with reduced induction of IL1-β and Tnf-α **(Fig. 1M, N, O).** Histopathology likewise showed preservation of colonic architecture in fasted-refed mice relative to continuously fed controls **(Supplementary Fig. 1H).**

Together, these findings indicate that short-term fasting purges pre-existing DNA-damaged IECs, thereby reducing susceptibility to inflammatory injury.

### Short-term fasting induces CDK5RAP3-dependent ER stress, driving apoptosis in DNA-damaged cells

To identify stress pathways engaged during fasting and linked to apoptotic susceptibility, we performed quantitative proteomic profiling of flow-purified IECs isolated from mice fasted for 12 h **(Fig. 2A)**. Fasting induced extensive proteome remodeling, with ∼1,007 proteins increased and 475 decreased at ≥1.5-fold **(Fig. 2B)**. Pathway enrichment analysis revealed a dominant endoplasmic reticulum (ER) stress signature together with activation of integrated stress response pathways **(Fig. 2C, D)**. ER stress emerged as one of the most strongly enriched programs, with coordinated induction of unfolded protein response (UPR) and ER-associated degradation (ERAD) components **(Fig. 2E)**.

**Figure 2.**
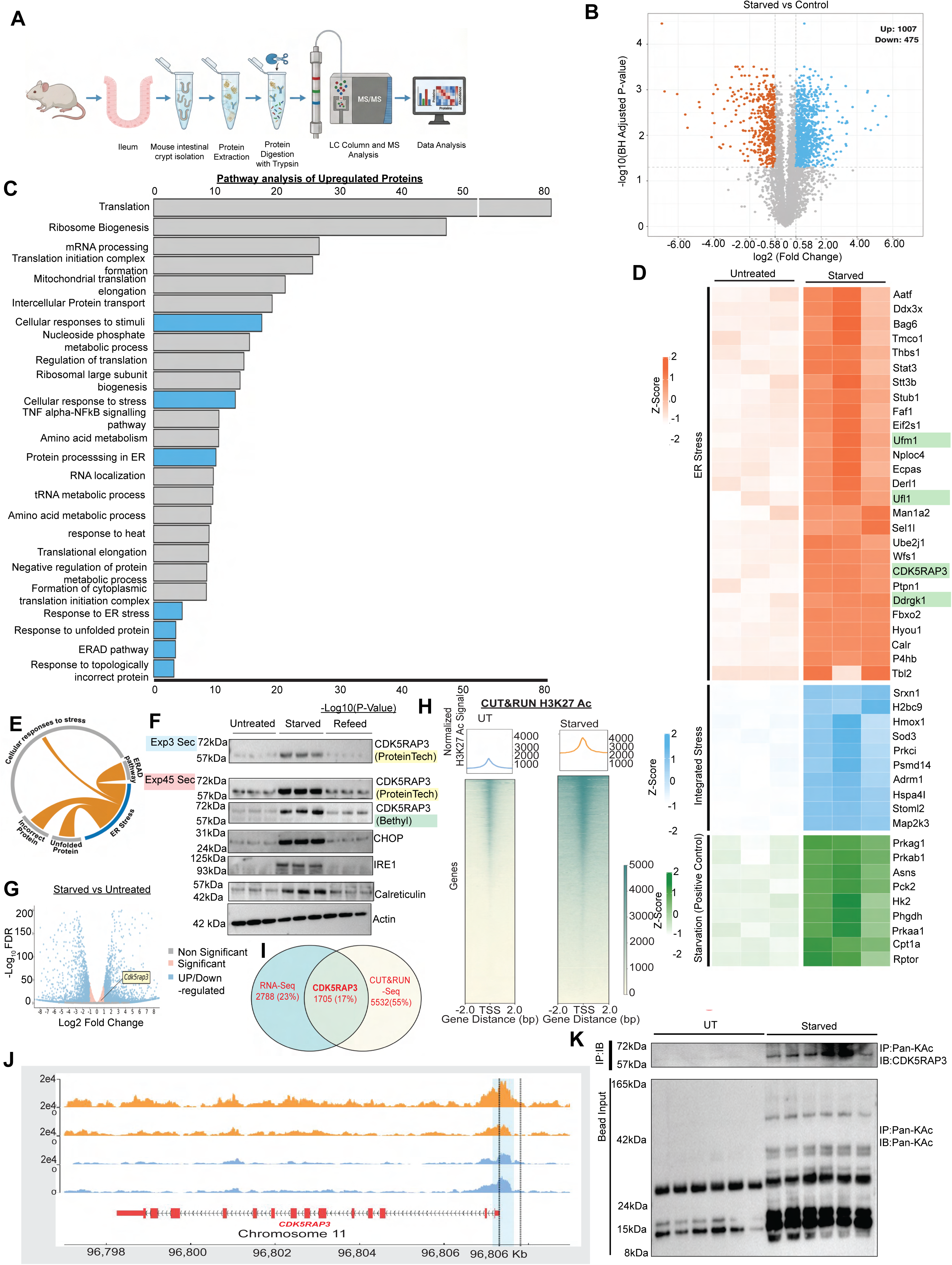
Fasting induces epigenetic and proteomic changes associated with elevated CDK5RAP3 and ER stress. **(A)** Workflow for proteomic analysis of FACS-purified IECs from Ileum. Schematic created in BioRender (CHATTERJEE, S., 2026;https://BioRender.com/mtrovit). **(B)** Volcano plot showing differentially expressed proteins in starved versus untreated mouse crypts. **(C)** Pathway enrichment analysis of upregulated proteins. **(D)** Heatmap of selected proteins associated with ER stress (red), integrated cellular stress (blue), and starvation responses (green, positive control), highlighting CDK5RAP3. **(E)** Network visualization of enriched biological processes and pathways related to cellular stress responses, including ER stress, unfolded protein response, and proteotoxic stress. **(F)** Immunoblot analysis of CDK5RAP3, CHOP, IRE1, and calreticulin in untreated, starved, and refeed conditions. The tests were conducted under two different exposures, 3 sec blue and 45 sec pink, and were validated using antibodies from Proteintech yellow and Bethyl green. **(G)** Volcano plot of bulk RNA sequencing from purified crypts of untreated and starved mice, highlighting CDK5RAP3 among differentially expressed transcripts during fasting. **(H)** Heat map of CUT&RUN analysis of H3K27ac signal in untreated and starved samples (highlighted blue). **(I)** Overlap of RNA-seq, and CUT&RUN datasets. **(J)** Genome browser tracks showing H3K27ac enrichment at the CDK5RAP3 transcription start sites (highlighted blue). **(K)** Immunoprecipitation of pan-acetylated proteins from 400 μg protein lysate followed by immunoblotting for CDK5RAP3 from untreated and starved samples. Data represent n = 3 independent biological replicates unless otherwise indicated; mean ± SD where applicable.

We validated ER stress response biochemically. Immunoblotting confirmed induction of calreticulin and robust upregulation of canonical ER stress markers, including CHOP and IRE1 **(Fig. 2F)**.

Because canonical ER stress markers largely report downstream signaling, we interrogated the fasting proteome to identify upstream control points. Among the most strongly induced proteins, components of the ribosome-associated UFMylation machinery, including Ufm1, Ufl1, Ddrgk1, and CDK5RAP3, emerged as prominent candidates **(Fig. 2D, F)**. This complex catalyzes UFM1 conjugation to the ribosomal protein RPL26 at the ER membrane, a modification essential for cotranslational proteostasis^17,18^. Consistently, RPL26 UFMylation supports ribosome-associated quality control, and its disruption is sufficient to activate ER stress signaling^19,20^.

Despite coordinated upregulation of Ufm1, Ufl1, CDK5RAP3, and Ddrgk1, this response could be functionally ineffective due to CDK5RAP3’s abbarent expression. CDK5RAP3 operates as a dosage-sensitive regulatory node: loss of CDK5RAP3 reduces RPL26 UFMylation^21^, whereas excess CDK5RAP3 paradoxically restricts UFM1 transfer and similarly diminishes RPL26 UFMylation^22^. Thus, fasting-driven accumulation of CDK5RAP3 could impedeRPL26 UFMylation despite increased expression of other core UFMylation machinery.

As expected, enforced CDK5RAP3 expression in cell sytem was sufficient to suppress RPL26 UFMylation and induce ER stress markers, including calreticulin, CHOP, and IRE1 **(Supplementary Fig. 2A)**. *In vivo*, fasting elicited a time-dependent increase in endogenous CDK5RAP3 beginning at 6 h and persisting through 12 h, coincident with loss of RPL26 UFMylation and subsequent induction of ER stress markers **(Supplementary Fig. 2B)**.

Notably, fasting-induced CDK5RAP3 expression and ER stress occurred to a similar extent in phospho-γ-H2AX⁺ and phospho-γ-H2AX⁻ IECs **(Supplementary Fig. 2C)**, indicating that ER stress is broadly induced across the epithelial compartment and is not restricted by baseline DNA damage status.

Next, to determine whether CDK5RAP3-driven ER stress cooperates with genomic instability to promote apoptosis. We overexpressed CDK5RAP3 in untreated cells and cells treated with UV radiation to create DNA damage. While CDK5RAP3 overexpression induced ER stress under both basal conditions and following UV-treatment; the cleaved caspase-3 was detected only under UV treated cells and was markedly enhanced by CDK5RAP3 overexpression **(Supplementary Fig. 2D)**. These results indicate that CDK5RAP3-driven ER stress broadly sensitizes cells but preferentially licenses apoptotic execution in the presence of DNA damage, recapitulating the selectivity observed *in vivo*.

Finally, this pathway is not restricted to the intestine. Fasting increased CDK5RAP3 abundance and ER stress markers in the liver, pancreas, and lung **(Supplementary Fig. 2E)**, indicating that the CDK5RAP3-ER stress axis operates across multiple tissues.

Together, fasting-induced CDK5RAP3 accumulation functionally disables ribosome-associated RPL26 UFMylation, thereby engaging ER stress. Although ER stress is broadly induced across the epithelium; apoptotic execution is selectively restricted to DNA-damaged cells.

### Fasting coordinates epigenetic and post-translational programs to increase CDK5RAP3 level

We next asked how fasting elevates CDK5RAP3 abundance. RNA-seq analysis of IECs from fed and fasted mice revealed increased Cdk5rap3 transcripts following fasting **(Fig. 2G)**, indicating a transcriptional component.

To define the chromatin basis of this response, we integrated RNA-seq with H3K27ac CUT&RUN profiling. Because H3K27ac marks active promoters and enhancers, we used it as a readout of fasting-induced epigenetic activation. Fasting elicited a global increase in H3K27ac across IEC chromatin **(Fig. 2H),** consistent with widespread transcriptional activation. Intersection of the RNA-seq and CUT&RUN datasets identified 1,705 upregulated genes that also gained promoter-associated H3K27ac **(Fig. 2I).** Cdk5rap3 was among these genes, indicating that its transcriptional induction occurs within a broader fasting-induced epigenetic program.

Locus-level analysis across a ±50 kb window surrounding the Cdk5rap3 transcription start site identified five significant fasting-induced H3K27ac peaks (FDR < 0.05), including one overlapping the TSS and four additional distal elements **(Fig. 2J; Supplementary Fig. 2F).** These data indicate coordinated activation of promoter-proximal and distal regulatory elements at the Cdk5rap3 locus during fasting.

However, the increase in Cdk5rap3 mRNA was modest relative to the marked accumulation of CDK5RAP3 protein, suggesting an additional post-translational layer of control. Because fasting broadly reshapes acetylation states and many chromatin-modifying enzymes also regulate non-histone substrates, we asked whether CDK5RAP3 itself is acetylated. CDK5RAP3 contains 35 lysine residues across its 530 amino acids, providing multiple candidate acetylation sites. Co-immunoprecipitation with a pan-acetyl-lysine antibody revealed minimal CDK5RAP3 acetylation in fed controls but robust acetylated CDK5RAP3 under fasting conditions **(Fig. 2K).**

Thus, fasting increases CDK5RAP3 levels through the coordinated action of two distinct, epigenetically-linked mechanisms: A) increased transcription driven by H3K27 acetylation at the promoter, and B) stabilization through increased protein-level acetylation.

### InsP₆ depletion during fasting inactivates HDAC3 and drives CDK5RAP3’s epigenetic and post-translational changes

To identify the upstream mechanism responsible for fasting-induced chromatin and CDK5RAP3 acetylation, we quantified histone acetyltransferase (HAT) and class I histone deacetylase (HDAC) activities in IECs. Fasting did not increase HAT activity **(Supplementary Fig. 3A),** but instead caused a profound reduction **(>90%)** in HDAC3 catalytic activity **(Fig. 3A)** without altering total HDAC3 protein abundance **(Fig. 3B).** Fasting also did not disrupt the association of HDAC3 with its obligate corepressor SMRT or compromise overall complex integrity **(Fig. 3C).** By contrast, the activities of other class I HDACs were largely preserved, with only a modest reduction in HDAC1 activity **(Fig. 3A).** HDAC3 is therefore the principal enzymatic target of fasting.

**Figure 3.**
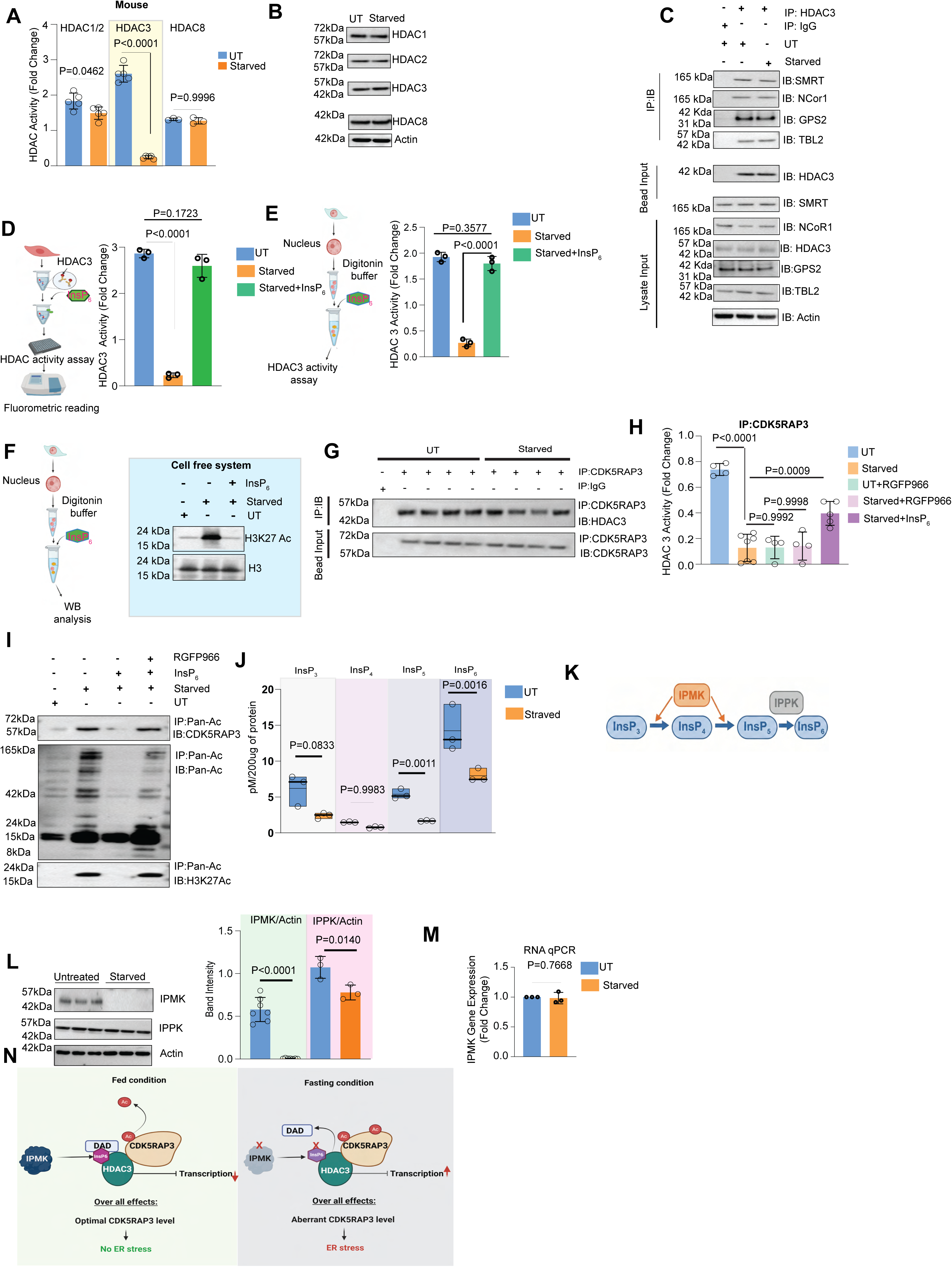
Fasting inactivates HDAC3 through depleting InsP₆. **(A)** Fluorometric class I HDAC activity assays from IECs from starved versus untreated (UT) mice (n ≥ 5; mean ± SD). **(B)** Immunoblot analysis from untreated (UT) and starved mice IECs (n=3). **(C)** Co-immunoprecipitation of HDAC3 or IgG from untreated (UT) and starved mouse IEC followed by immunoblotting with SMRT, NCoR1, GPS2, and TBL2. (n=3). **(D)** Schematics of *in vitro* InsP_6_ treatment and HDAC3 activity assay *in vitro* (Created in BioRender. CHATTERJEE, S. (2026) https://BioRender.com/mtrovit). HDAC3 immunopurified from IECs of untreated (UT) or starved mice. InsP₆ (100 nM) was treated prior to the HDAC3 activity assay. (n=3;mean ± SD). **(E)** A schematic of InsP_6_ treatment followed by HDAC3 activity assay in a cell-free nuclear system (Created in BioRender. CHATTERJEE, S. (2026) https://BioRender.com/mtrovit). Nuclei from IECs of starved mice were permeabilized and treated with 1µM InsP₆ before HDAC3 activity measurement (n=3;mean ± SD). **(F)** A schematic (Created in BioRender. CHATTERJEE, S. (2026) https://BioRender.com/mtrovit) of InsP₆ treatment in a cell-free nuclear system followed by western blot of H3k27ac. (n=3). **(G)** Anti-CDK5RAP3 immunoprecipitation from 400μg followed by HDAC3 immunoblotting shows interaction between CDK5RAP3 and HDAC3 (n = 4). **(H)** CDK5RAP3-bound HDAC3 pool was purified from 400 μg of protein, followed by HDAC3 activity assay. RGFP966 (HDAC3 inhibitor) and InsP₆ (100 nM) were treated *in vitro* in the reaction (n ≥ 4; mean ± SD). **(I)** Pan-acetylation immunoprecipitation from 400 μg of protein followed by CDK5RAP3 immunoblotting reveals increased CDK5RAP3 acetylation under GFD, rescued by *in vitro* InsP₆ treatment. (n = 3). **(J)** Mass spectrometry–based quantification of higher-order inositol phosphates in IECs demonstrates. (n=3;mean ± SD) **(K)** Schematic of the higher-order inositol phosphate (HOIP) biosynthetic pathway (Created in BioRender. CHATTERJEE, S. (2026) https://BioRender.com/mtrovit). **(L)** Immunoblot followed by densitometric analysis of IPMK and IPPK from mouse IECs. (IPMK n=7, IPPK n=3;mean ± SD). **(M)** Quantitative RT–PCR of *Ipmk* transcript levels in IECs from untreated (UT) and starved mice. (n=3;mean ± SD). **(N)** Schematics depicting mechanistic hypotheses.

Our previous work^10,23^ established that InsP_6_ is an essential cofactor for HDAC3 catalytic activity. Loss of InsP_6_ inactivates HDAC3 without altering HDAC3 protein abundance or abolishing corepressor assembly, in part by weakening the interaction between HDAC3 and the SMRT deacetylase-activating domain (DAD). Consistent with this mechanism, HDAC3 purified from IECs of fasted mice rapidly and fully regained activity after brief incubation with 0.1 μM InsP_6_ **(Fig. 3D),** demonstrating a direct requirement for InsP_6_ in maintaining HDAC3 catalysis during fasting.

We next tested this in a complementary cell-free nuclear system. Addition of InsP_6_ to nuclei isolated from fasted IECs robustly restored HDAC3 activity **(Fig. 3E)** and reset global H3K27 acetylation to levels observed in non-fasted controls **(Fig. 3F).** These findings establish that InsP_6_ depletion is sufficient to account for the loss of HDAC3 activity and the resulting chromatin hyper-acetylation phenotype.

Because fasting increased CDK5RAP3 acetylation, we asked whether HDAC3 directly regulates CDK5RAP3 deacetylation. Endogenous CDK5RAP3 immuno-precipitates from fed and fasted IECs retained HDAC3 association under both conditions **(Fig. 3G)**, indicating that complex assembly is preserved. However, HDAC3 co-purifying with CDK5RAP3 from fasted IECs exhibited markedly reduced deacetylase activity compared to HDAC3 recovered from fed controls **(Fig. 3H)**, indicating catalytic inactivation without loss of interaction.

To enable quantitative comparison, CDK5RAP3 immunoprecipitation conditions were adjusted to normalize protein recovery between fed and fasted samples **(Supplementary Figure 3B)**, accounting for fasting-induced increases in CDK5RAP3 abundance. Under these conditions, deacetylase activity measured in CDK5RAP3 immunoprecipitates was lower than that obtained by direct HDAC3 immunoprecipitation **(Fig. 3A)**, consistent with selective assessment of the CDK5RAP3-bound pool. This activity was abolished by the HDAC3-selective inhibitor RGFP966 **(Fig. 3H)**, confirming specificity. Notably, addition of InsP_6_ *in vitro* restored deacetylase activity to CDK5RAP3-associated HDAC3 was purified from fasted IECs **(Fig. 3H)** and concomitantly reduced CDK5RAP3 acetylation **(Fig. 3I),** indicating that HDAC3-dependent deacetylation of CDK5RAP3 is InsP_6_-sensitive.

Consistent with this mechanism, mass spectrometry-based profiling revealed a significant reduction in InsP_6_ in IECs from fasted mice **(Fig. 3J)**. These data position InsP₆ production as the metabolic input that sustains HDAC3 catalytic activity. We therefore examined the inositol phosphate biosynthetic pathway **(Fig. 3K)**. Among enzymes required for intracellular InsP_6_ synthesis, fasting selectively reduced inositol polyphosphate multikinase (IPMK) protein abundance, whereas inositol-pentakisphosphate 2-kinase (IPPK) remained unchanged **(Fig. 3L)**. Notably, Ipmk mRNA levels were unaffected **(Fig. 3M)**, indicating post-transcriptional destabilization of IPMK.

Together, these data identify InsP₆ depletion as the upstream event that inactivates HDAC3 during fasting. This loss of HDAC3 activity leads to an increase in both H3K27 acetylation on the Cdk5rap3 promoter and the post-translational acetylation of the CDK5RAP3 protein itself. Although HDAC3 remains physically associated with CDK5RAP3 even during fasting, its catalytic activity is impaired, resulting in reduced deacetylation of CDK5RAP3. These coordinated epigenetic and post-translational changes drive CDK5RAP3 accumulation **(Fig. 3N)**.

### Cell-based screen identifies that growth factor deprivation induces CDK5RAP3 mediated ER stress and phenocopies fasting-associated mechanisms

Having defined how fasting elevates CDK5RAP3, we next asked what upstream signal disrupts the InsP₆-HDAC3 axis. To dissect, we used genetically tractable cell systems, which permit precise and temporally controlled perturbations that are difficult to achieve *in vivo*. Using these cell-based assays, we defined the detailed mechanism **(Figs. 4–6)** and then tested its physiological relevance by confirming key pathway nodes in mouse intestine and enteroids **(Fig. 7)**, demonstrating that the mechanisms uncovered in cells operate in tissue.

**Figure 4.**
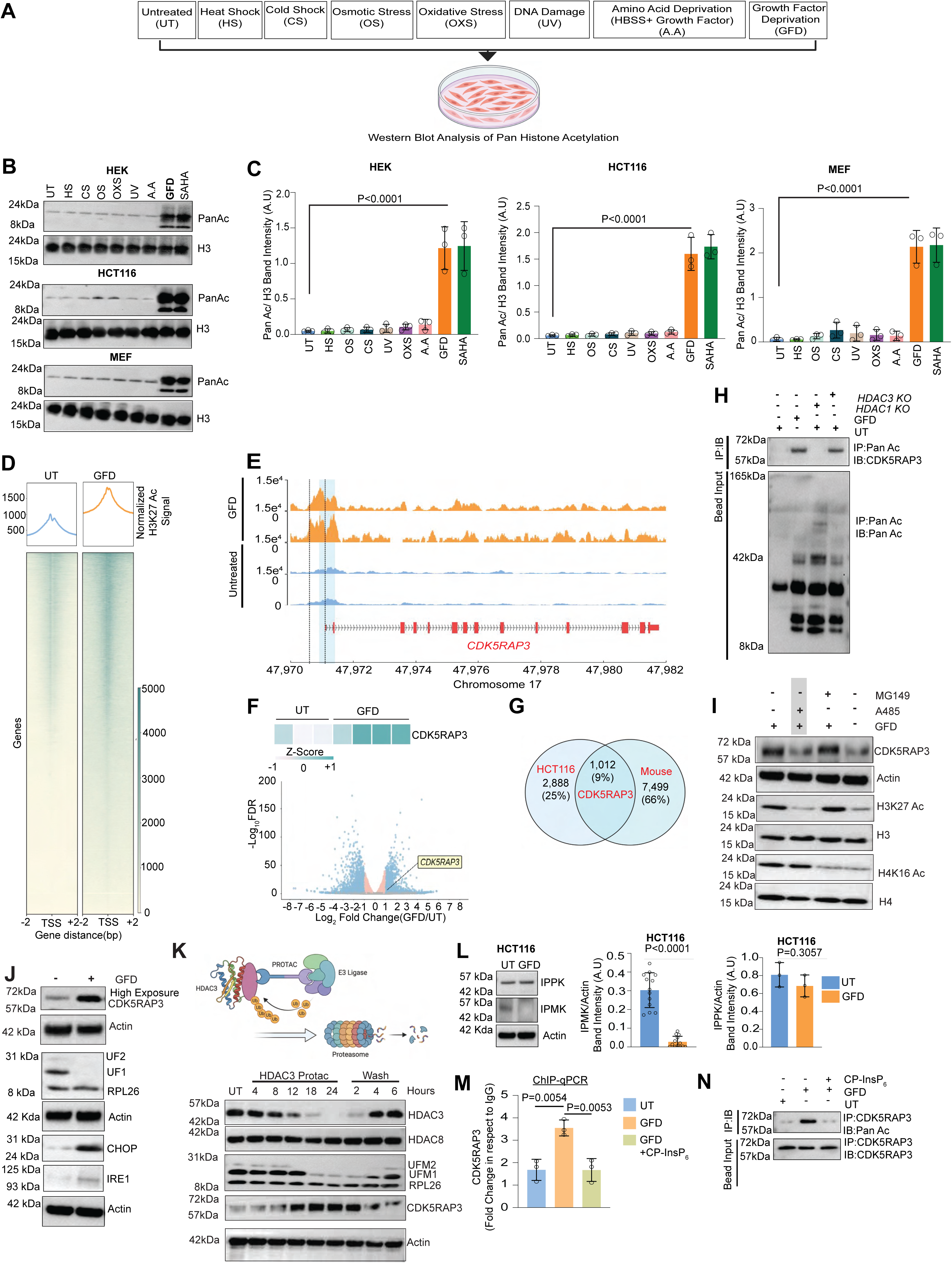
Growth factor deprivation mimics fasting-induced chromatin remodeling and CDK5RAP3 gene expression. **(A)** Experimental conditions including untreated (UT), heat shock (HS), cold shock (CS), osmotic stress (OS), oxidative stress (OXS), UV-induced DNA damage (UV), amino acid deprivation (A.A), and growth factor deprivation (GFD) followed by western blot. Created in BioRender. CHATTERJEE, S. (2026) https://BioRender.com/mtrovit. **(B)** Immunoblot analysis of pan-histone acetylation (PanAc) and total H3 in HEK, HCT116, and MEF cells under the indicated conditions. **(C)** Quantification of PanAc normalized to H3 in HEK, HCT116, and MEF cells (n = 3; mean ± SD). **(D)** Heat map of CUT&RUN analysis showing H3K27ac signal distribution in UT and GFD conditions in HCT116 cells. (n = 2). **(E)** Genome browser tracks showing H3K27ac enrichment at the CDK5RAP3 locus in untreated and GFD HCT116 cells. (n = 2). **(F)** Heatmap and volcano plot of gene expression changes under GFD in HCT116 cells highlighting CDK5RAP3. **(G)** Overlap of CDK5RAP3-associated datasets from HCT116 and mouse samples. **(H)** Immunoprecipitation of acetylated proteins (from 400 μg lysate) followed by immunoblotting for CDK5RAP3 from HDAC1 and HDAC3 knockout HCT116 cells under UT and GFD conditions. (n=3). **(I)** Immunoblot analysis of CDK5RAP3, H3K27ac, H4K16ac, and histone controls following treatment with histone acetyltransferase inhibitors under GFD conditions. MG149 (35µM) Tip60 inhibitor, A485 (10µM) P300 inhibitor treated in GFD for 12 hours.(n=3) **(J)** Immunoblot analysis of CDK5RAP3, RPL26 UFMylation, CHOP, and IRE1 in UT and GFD HCT116 cells.(n=3) **(K)** Schematic of HDAC3 PROTAC-mediated degradation. Created in BioRender (CHATTERJEE, S., 2026; https://BioRender.com). Western blot analysis in HCT116 cells assessed the time-dependent degradation of HDAC3 after PROTAC treatment and the effect of compound washout. **(L)** Immunoblot and quantification of IPMK and IPPK protein levels in HCT116 cells (n = 3 to 13; mean ± SD). **(M)** ChIP–qPCR analysis of CDK5RAP3 under UT, GFD, and GFD+CP-InsP₆ treated HCT116 cells (n = 3; mean ± SD). **(N)** Immunoprecipitation of CDK5RAP3 from 400 μg of protein lysate from HCT116 cells followed by immunoblotting for acetylation under indicated conditions. (n = 3) All dots in bar plots represent individual biological replicates.

**Figure 5:**
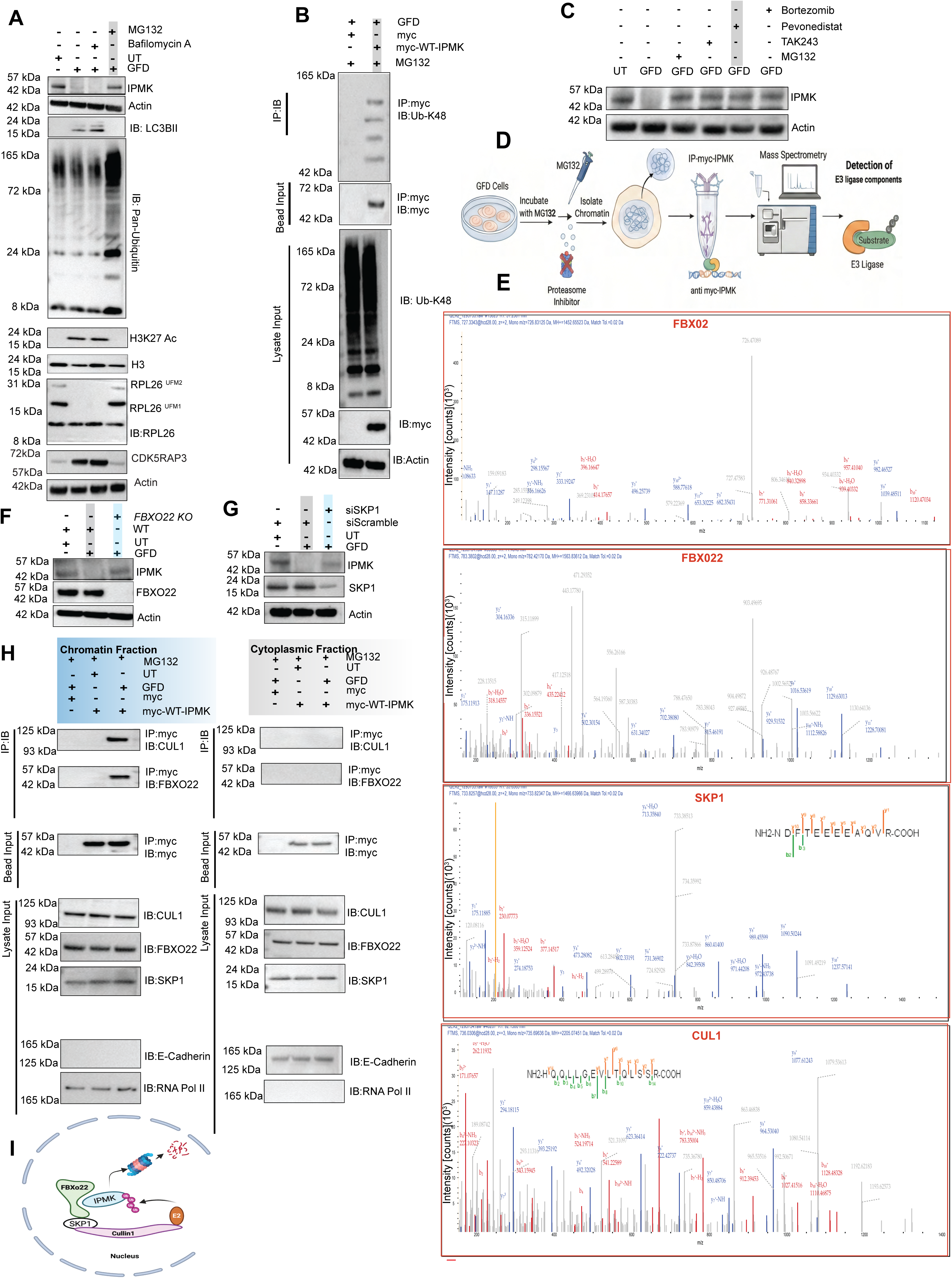
FBXO22–SCF–Cullin1 complex mediates proteasomal degradation of nuclear IPMK during growth factor deprivation. **(A)** Immunoblot analysis of IPMK, H3K27Ac, CDK5RAP3, polyubiquitin, and LC3B in untreated and GFD-treated cells with MG132 or bafilomycin A1; actin serves as a loading control **(n = 3)**. **(B)** Immunoprecipitation of WT-myc-IPMK followed by immunoblotting for K48-linked ubiquitin in untreated and GFD conditions in the presence of MG132 (proteasome inhibitor) (n = 3). **(C)** Immunoblot analysis of IPMK levels in GFD-treated cells following treatment with TAK-243, pevonedistat, MG132, or bortezomib (n = 3). **(D)** Schematic of experimental workflow for identification of IPMK-associated ubiquitin ligase components. Chromatin-associated myc-IPMK complexes were isolated from MG132-treated GFD cells and analyzed by mass spectrometry. Created with BioRender; Chatterjee, S., 2026; https://BioRender.com/mtrovit. **(E)** Mass spectrometry spectra for Cullin1, SKP1, FBXO2, and FBXO22 as IPMK-associated complexes (n = 3). **(F)** Immunoblot analysis of IPMK in A549 cells following FBXO22 depletion under indicated conditions (n = 3). **(G)** Immunoblot analysis of IPMK following SKP1 knockdown in untreated and GFD HCT116 cells (n = 3). **(H)** Subcellular fractionation and co-immunoprecipitation of IPMK with FBXO22, SKP1, and Cullin1 in chromatin and cytoplasmic fractions under indicated conditions (n = 3). **(I)** Model of IPMK ubiquitination by an FBXO22–SKP1–Cullin1–RBX1 complex.

**Figure 6.**
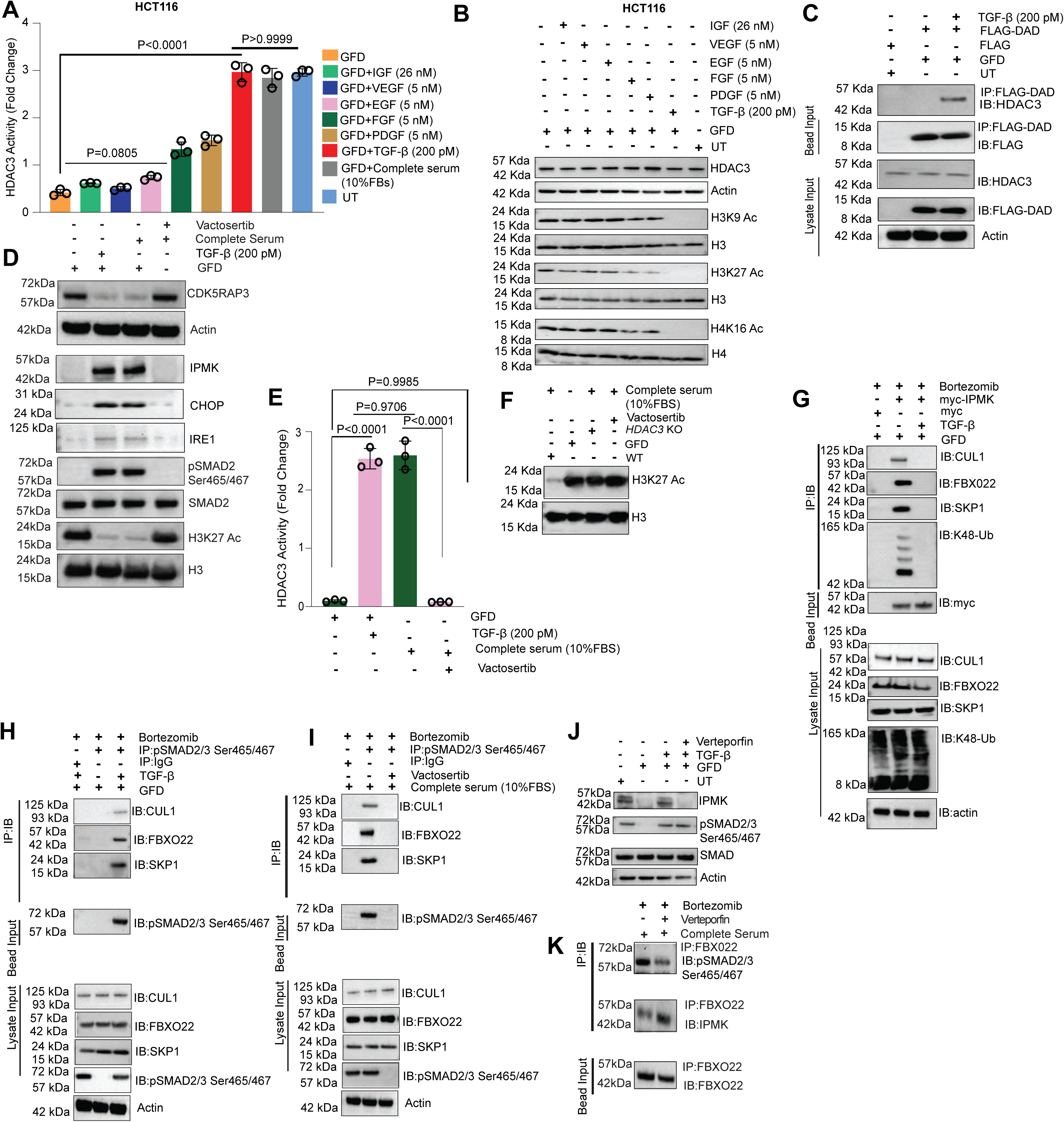
TGFβ signaling stabilizes the IPMK–HDAC3 axis by preventing FBXO22–SCF–Cullin1–mediated IPMK ubiquitination. **(A)** HDAC3 activity assay in GFD-treated HCT116 cells following stimulation (3 h) with TGFβ, IGF1, VEGF, EGF, FGF, or PDGF. (n = 3; mean ± SD). **(B)** Immunoblot analysis of H3K9ac, H3K27ac, H4K16ac, and HDAC3 in untreated, GFD, and growth factor-stimulated conditions. (n = 3). **(C)** Co-immunoprecipitation of HDAC3 from HCT116 cells overexpressing FLAG-SMRT deacetylase-activating domain (FLAG-DAD) under indicated conditions. (n = 3). **(D)** Immunoblot analysis of IPMK, CDK5RAP3, CHOP, IRE1, and H3K27ac following treatment with the TGFβ receptor inhibitor vactosertib in HCT116 cells. (n = 3). **(E)** HDAC3 enzymatic activity assay in HCT116 cells treated with TGF-β, and vactosertib. (n = 3; mean ± SD). **(F)** Immunoblot analysis of H3K27ac in untreated, GFD, vactosertib-treated, and HDAC3 knockout cells. (n = 3). **(G)** Immunoprecipitation of IPMK–myc followed by immunoblotting for FBXO22, SKP1, Cullin1, and K48-linked ubiquitin under the indicated conditions. (n = 3). **(H)** Co-immunoprecipitation of phosphorylated SMAD2/3 with FBXO22, SKP1, and Cullin1 under the indicated conditions. (n = 3). **(I)** Co-immunoprecipitation of phosphorylated SMAD2/3 with FBXO22–SCF–Cullin1 components following vactosertib treatment. (n = 3). **(J)** IPMK expression and pSMAD2/3 level after Verteportin (blocks pSmad2/3 nuclear localization) treatment. (n = 3). **(K)** Immunoprecipitation study to show effects of Verteportin on FBXO22 interaction with IPMK and pSMAD2/3. (n = 3).

**Figure 7.**
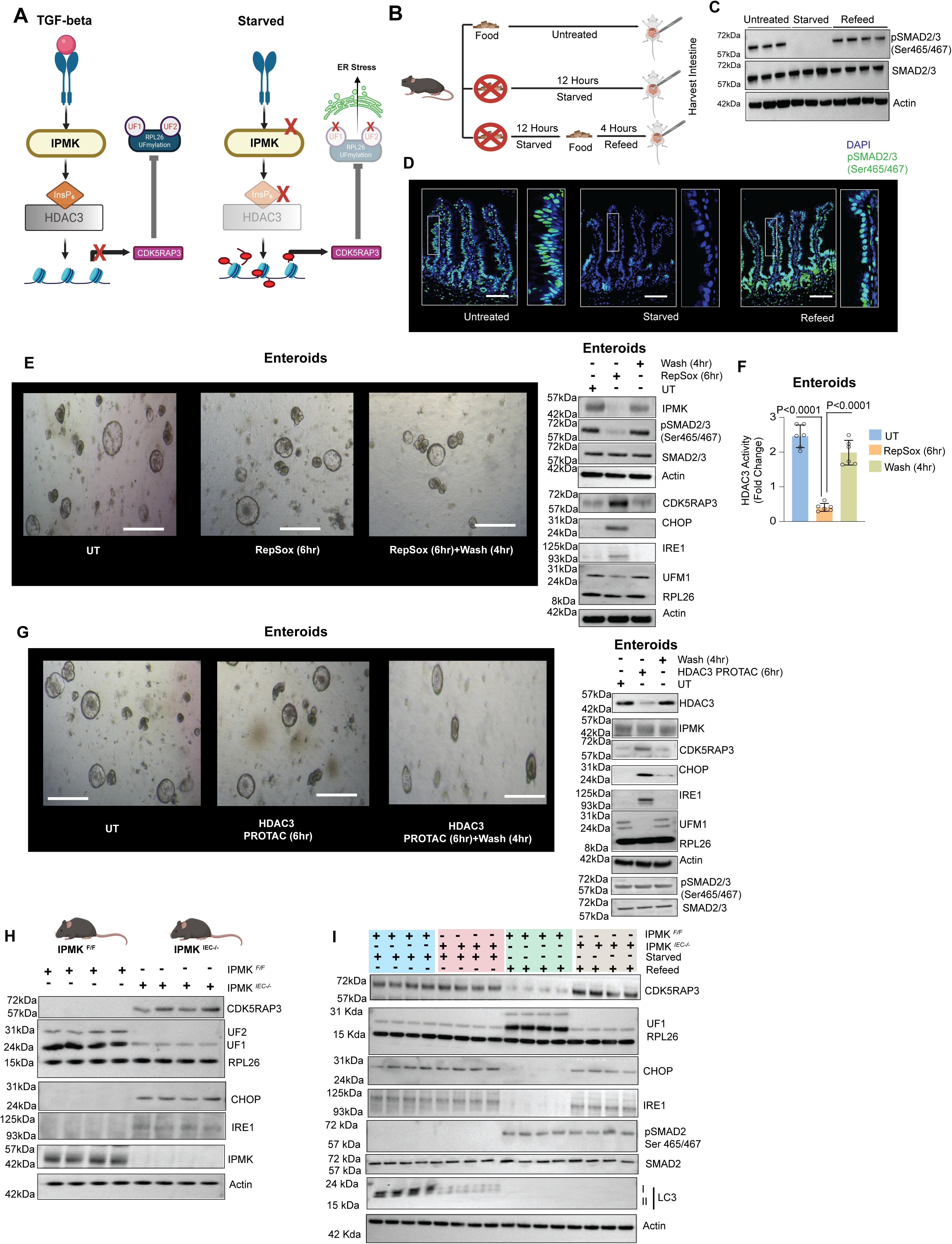
Fasting suppresses TGF-β signaling and destabilizes the IPMK–HDAC3 axis to induce ER stress in intestinal epithelium. **(A)** Schematic illustration. Created with BioRender; Chatterjee, S., 2026; https://BioRender.com/mtrovit. **(B)** Experimental schematic of fasting and refeeding protocol used in mice. Created with BioRender; Chatterjee, S., 2026; https://BioRender.com/mtrovit. **(C)** Immunoblot analysis of ileal crypts showing phosphorylated pSMAD2/3 (Ser465/467) in untreated (UT), starved (12 h), and refed (24h) conditions (UT n = 3, starved n = 3, refed n = 4). **(D)** Immunofluorescence staining of ileal epithelium showing pSMAD2/3 localization under indicated conditions; scale bar, 50 μm. (UT n = 3, starved n = 3, refed n = 4). **(E)** Representative bright-field images of intestinal enteroids. Scale bar 60µm. Enetroids cultured from wild-type mice were treated with RepSox on day 3 (n = 3 mice). Organoids were also tested by Western blotting after washing off RepSox with normal media. **(F)** HDAC3 activity assay tested from enteroids. (n = 3;mean ± SD) **(G)** Representative bright-field images of intestinal enteroids. Scale bar 60µm. Enteroids treated with an HDAC3 PROTAC on day 3, followed by western blot. (n = 3 mice). Organoids were also tested using Western blots after washing off the PROTAC with normal media. **(H)** Immunoblot analysis from ileul crypts of untreated IPMK floxed (*IPMK^F/F^*) and intestine-specific IPMK-depleted line (*IPMK^IEC-/^*^-^)(n = 4 mice). **(I)** Immunoblot analysis from ileal crypts of untreated IPMK floxed (*IPMK^F/F^*) and intestine-specific IPMK-depleted line (*IPMK^IEC-/-^*) after fasting and refeeding. (n = 4 mice).

We first asked which fasting-associated signal drives the epigenetic remodeling observed in mouse IECs. Immortalized cell lines were exposed for 12 h either to amino acid starvation or to complete growth factor deprivation (GFD). GFD induced a robust global increase in histone acetylation comparable to that triggered by the pan-HDAC inhibitor SAHA, whereas amino acid starvation had minimal effect **(Fig. 4A-C).** Other stress paradigms, including oxidative stress, UV irradiation, and temperature shock, failed to reproduce this signature **(Fig. 4A-C).** Thus, histone hyperacetylation arises specifically from loss of growth factor signaling.

To define the chromatin basis of this response, we combined quantitative SILAC proteomics **(Supplementary Fig. 4A-C)** with CUT&RUN profiling **(Fig. 4D)** of HCT116 cells. Both approaches converged on H3K27 acetylation as the dominant chromatin modification induced by GFD. Genome-wide mapping revealed a global increase in H3K27ac **(Fig. 4D)** with strong enrichment at the CDK5RAP3 promoter **(Fig. 4E),** closely resembling the chromatin landscape observed in fasted IECs **(Fig. 2H, J).** Consistent with this remodeling, RNA sequencing demonstrated induction of CDK5RAP3 **(Fig. 4F).** Cross-species comparison identified 1,012 genes shared between GFD-treated HCT116 cells and fasted mouse IECs **(Fig. 4G),** indicating that GFD captures a conserved fasting-induced transcriptional program *in vitro*.

Because CDK5RAP3 emerged as a prominent shared target, we next examined how its expression is regulated. In addition to transcriptional induction, GFD increased CDK5RAP3 acetylation **(Fig. 4H),** paralleling the fasting response *in vivo* **(Fig. 2K).** Genetic depletion showed that loss of HDAC3, but not HDAC1, was sufficient to induce CDK5RAP3 acetylation **(Fig. 4H),** indicating selective regulation by HDAC3. Immunoprecipitation further revealed direct interaction between HDAC3 and CDK5RAP3 **(Supplementary Fig. 4D).** CDK5RAP3 also associated with the histone acetyltransferase p300 **(Supplementary Fig. 4E)**, suggesting opposing enzymatic control. Consistent with this model, inhibition of p300 with A485 attenuated CDK5RAP3 induction during GFD **(Fig. 4I),** whereas inhibition of Tip60 with MG149 had little effect. Thus, p300-dependent acetylation and HDAC3-mediated deacetylation coordinately regulate CDK5RAP3 expression and modification.

If CDK5RAP3 accumulation is functionally relevant, it should affect downstream proteostasis pathways. Indeed, GFD markedly increased CDK5RAP3 protein levels, reduced RPL26 UFMylation, and induced ER stress markers, including CHOP and IRE1 **(Fig. 4J).** These changes phenocopied the proteostatic defects observed in IECs during fasting **(Fig. 2F),** positioning CDK5RAP3 accumulation as a central driver of ER stress.

Because fasting selectively suppresses HDAC3 activity *in vivo* **(Fig. 3A),** we next asked whether GFD targets this enzyme. GFD selectively reduced HDAC3 catalytic activity **(Supplementary Fig. 4F, G),** closely paralleling the reduction observed in fasted IECs **(Fig. 3A).** This effect was conserved across eight mammalian cell lines **(Supplementary Fig. 4H, I),** indicating a broadly conserved growth factor-dependent regulatory mechanism.

Genetic disruption of HDAC3 derepressed CDK5RAP3 expression **(Supplementary Fig. 4J),** reduced RPL26 UFMylation, and induced ER stress markers. Acute HDAC3 degradation using an HDAC3-directed PROTAC produced the same phenotype **(Fig. 4K).**

We next examined how HDAC3 activity is suppressed. Expression of wild-type HDAC3 restored repression of CDK5RAP3 **(Supplementary Fig. 4K),** whereas an InsP₆-binding-defective mutant (HDAC3^HOIP^) failed to rescue these defects **(Supplementary Fig. 4K),** demonstrating a requirement for InsP₆-dependent HDAC3 catalytic activity. Consistent with this, GFD did not disrupt the HDAC3-SMRT interaction **(Supplementary Fig. 4L)** but selectively weakened association with the SMRT deacetylase-activating domain (DAD), which is required for HDAC3 activation **(Supplementary Fig. 4M).** Importantly, this defect was fully restored by addition of cell-permeable InsP₆ **(Supplementary Fig. 4M),** indicating that InsP₆ depletion destabilizes the active HDAC3-DAD complex without disrupting the core HDAC3-SMRT scaffold.

Because HDAC3 activity depends on InsP₆ availability, we next examined the InsP₆ biosynthetic pathway. Growth factor deprivation selectively reduced IPMK protein abundance **(Fig. 4L)** without affecting IPPK levels, resulting in decreased intracellular InsP₆ **(Supplementary Fig. 4N)** despite unchanged IPMK mRNA **(Supplementary Fig. 4O).** This mirrors the post-transcriptional loss of IPMK observed in fasted IECs.

Consistent with this mechanism, addition of cell-permeable InsP₆ (CP-InsP₆) restored chromatin homeostasis. CP-InsP₆ reduced H3K27ac at the CDK5RAP3 locus **(Fig. 4M),** decreased CDK5RAP3 acetylation **(Fig. 4N), a**nd normalized CDK5RAP3 expression, thereby restoring RPL26 UFMylation and suppressing ER stress markers **(Supplementary Fig. 4P).**

Because CDK5RAP3 is an essential scaffold for the UFMylation machinery^24–27^, its genetic depletion also diminishes RPL26 UFMylation and triggers ER stress^21^.Conversely, aberrant CDK5RAP3 accumulation prevents UFM transfer to RPL26 and likewise impairs UFMylation^22^, precluding a clear rescue strategy based on CDK5RAP3’s genetic depletion.

These data collectively indicate that the effects induced by fasting are clearly mimicked in a cell-based system. An unbiased screen of various nutrient and metabolic stress signals revealed that only growth factor deprivation (GFD) increased the level of CDK5RAP3 through both transcriptional and post-translational changes. Intriguingly, although GFD disrupts the IPMK-HDAC3 epigenetic axis by decreasing IPMK protein levels, the overall ER stress and CDK5RAP3 level were successfully restored to normal simply by treating the cells with CP-InsP_6_.

### Growth factor deprivation depletes InsP₆ via proteasomal degradation of nuclear IPMK

Because GFD reduced IPMK protein abundance without impacting transcription, we next asked how IPMK is destabilized. Proteasome inhibition with MG132, but not autophagy inhibition with bafilomycin A1, fully restored IPMK levels in GFD-treated cells **(Fig. 5A).** MG132 also normalized global H3K27 acetylation and suppressed induction of CDK5RAP3 **(Fig. 5A),** indicating that GFD engages an ubiquitin-proteasome pathway upstream of chromatin remodeling and ER stress signaling.

Consistent with this, myc-IPMK immunoprecipitation followed by ubiquitin immunoblotting revealed strong K48-linked polyubiquitination of IPMK under GFD **(Fig. 5B),** demonstrating that GFD promotes IPMK ubiquitination and proteasomal degradation. Inhibition of ubiquitin activation with the E1 inhibitor TAK-243 or blockade of Cullin1 neddylation (required for optimal Cullin1-based SCF complex function) with pevonedistat likewise preserved IPMK levels **(Fig. 5C),** implicating a Cullin1-based SCF E3 ligase in IPMK turnover.

To identify the substrate adaptor responsible for IPMK recruitment, we enriched transient E3-substrate intermediates by treating GFD cells with MG132 and subjected chromatin-associated IPMK complexes to mass spectrometry **(Fig. 5D).** This identified the core SCF components Cullin1 and SKP1 together with two candidate F-box proteins, FBXO2 and FBXO22, as IPMK-interacting partners **(Fig. 5E).** Genetic depletion of FBXO22 **(Fig. 5F),** but not FBXO2 **(Supplementary Fig. 5A),** completely prevented IPMK loss during GFD, identifying FBXO22 as the principal substrate-recognition module mediating IPMK ubiquitination. Depletion of *SKP1*, a core SCF scaffold, similarly preserved IPMK levels during GFD **(Fig. 5G).**

Subcellular fractionation revealed that this interaction is spatially restricted. Nuclear IPMK selectively co-purified with FBXO22 and Cullin1 in GFD-treated cells, whereas these interactions were absent in serum-fed controls and undetectable in the cytoplasmic IPMK pool under either condition **(Fig. 5H).** These data indicate that GFD selectively targets nuclear IPMK for recruitment to an FBXO22-SKP1-Cullin1 SCF ligase, resulting in K48-linked polyubiquitination and proteasomal degradation **(Fig. 5I).**

Thus, growth factor deprivation triggers FBXO22–Cullin1-dependent degradation of nuclear IPMK. Because IPMK is predominantly nuclear^28^, its loss depletes nuclear InsP₆, thereby inactivating HDAC3. Notably, whereas the initiating metabolic signal originates in the nucleus through IPMK loss, the CDK5RAP3 protein primarily localizes to the ER^29^, suggesting that nuclear InsP₆ licenses HDAC3 catalytic activity before its redistribution to the cytosol, whereas depletion of nuclear InsP₆ during growth factor deprivation prevents this activation, leaving CDK5RAP3-associated HDAC3 catalytically inactive and thereby promoting CDK5RAP3 stabilization.

Collectively, GFD engages FBXO22-Cullin1-dependent E3 ubiquitin ligase degrading IPMK.

### CDK5RAP3 acetylation under GFD prevents its autophagic degradation, leading to its accumulation

Next, after determining the mechanism by which GFD triggers the proteasomal degradation of IPMK, we investigated how the CDK5RAP3’s post-translational modification, mediated by acetylation, influences its protein stability and accumulation.

Under basal conditions, inhibition of autophagic flux with bafilomycin A1 resulted in robust accumulation of CDK5RAP3, whereas proteasome inhibition with bortezomib had minimal effect **(Supplementary Fig. 6A)**. Consistently, genetic or pharmacologic disruption of autophagy, using *ATG5* KO cells or the ULK1/2 inhibitor SBI-0206965, was sufficient to drive CDK5RAP3 accumulation **(Supplementary Fig. 6C)**, indicating that CDK5RAP3 is constitutively degraded via the autophagic pathway.

We asked whether CDK5RAP3 stabilization during GFD reflects impaired autophagic degradation. Suppression of CDK5RAP3 acetylation, either by p300 inhibition with A485 or by supplementation with cell-permeable InsP₆, prevented CDK5RAP3 accumulation during GFD **(Supplementary Fig. 6D)**. In contrast, blockade of autophagic flux with bafilomycin A1 restored CDK5RAP3 accumulation under all conditions, including when acetylation was suppressed **(Supplementary Fig. 6D)**.

Together, these findings establish that CDK5RAP3 stability is governed by autophagic turnover and that GFD-induced acetylation stabilizes CDK5RAP3 by limiting its autophagic degradation.

### TGF-β signaling maintains the IPMK-HDAC3 axis to suppress ER stress and CDK5RAP3 accumulation

We next sought to identify the specific growth factor pathway responsible for maintaining the IPMK-HDAC3 epigenetic axis, given that GFD disrupts this axis and consequently elevates CDK5RAP3-driven ER stress.

A screen of growth factors revealed that TGFβ is the dominant regulator capable of restoring both IPMK protein levels and the enzymatic activity of HDAC3 **(Fig 6A-D)**. Notably, this effect was observed even at a low concentration of 200 picomolar after just 3 hours of treatment. In detail, TGFβ restored HDAC3 enzymatic activity **(Fig. 6A, Supplementary Fig. 7A)** and normalized global histone acetylation to near-basal levels across multiple cell lines **(Fig. 6B, Supplementary Fig. 7B).** PDGF and FGF produced modest effects, whereas IGF1, VEGF, and EGF failed to elicit detectable responses. Mechanistically, TGFβ stimulation restored the interaction between HDAC3 and the SMRT deacetylase-activating domain (DAD) **(Fig. 6C),** which is disrupted upon GFD.

Importantly, TGFβ treatment restored IPMK protein to levels comparable to complete serum, while CDK5RAP3 and ER stress markers was markedly reduced **(Fig. 6D)**. The specificity of TGFβ signaling was confirmed by pharmacologic inhibition with the type I receptor inhibitors vactosertib or RepSox in serum-fed cells, which caused pronounced IPMK loss together with robust CDK5RAP3 induction **(Fig. 6D, Supplementary Fig. 7C)**. TGFβ blockade also increased CHOP and IRE1 **(Fig. 6D)**. Consistent with these effects, vactosertib reduced HDAC3 activity and elevated global histone acetylation **(Fig. 6E, F),** fully recapitulating the GFD state.

Notably, a growth factor cocktail containing IGF1, VEGF, EGF, FGF, and PDGF, but lacking TGFβ, failed to activate HDAC3 **(Supplementary Fig. 7E)** or restore IPMK and CDK5RAP3 expression **(Supplementary Fig. 7F),** underscoring the specificity of the TGFβ pathway.

Because IPMK degradation during GFD occurs through FBXO22-Cullin1-mediated ubiquitination, we next asked whether TGFβ regulates this step. Immunoprecipitation analyses revealed that TGFβ stimulation disrupted the interaction between IPMK and FBXO22 **(Fig. 6G),** thereby dissociating IPMK from the FBXO22-Cullin1 complex. Consistently, TGFβ markedly reduced K48-linked ubiquitination of IPMK **(Fig. 6G),** indicating that TGFβ stabilizes IPMK by suppressing its ubiquitination and proteasomal degradation.

We next investigated how TGFβ signaling displaces IPMK from the FBXO22-SCF complex. Because TGFβ activation promotes nuclear accumulation of phosphorylated SMAD2/3 (pSMAD2/3), which themselves undergo ubiquitin-dependent turnover^30^, we hypothesized that pSMAD2/3 compete with IPMK for FBXO22 binding. Consistent with this model, chromatin fractionation and co-immunoprecipitation showed that pSMAD2/3 associated with FBXO22 and the SCF-Cullin1 complex following TGFβ stimulation **(Fig. 6H),** whereas this interaction was lost during GFD or pharmacological inhibition of TGFβ signaling **(Fig. 6I).**

To determine whether nuclear pSMAD2/3 is required, we treated cells with verteporfin, which disrupts the pSMAD2/3-SMAD4 complex and prevents nuclear localization of pSMAD2/3 without affecting SMAD2/3 phosphorylation^31^. Verteporfin did not alter SMAD2/3 phosphorylation but abolished the ability of TGFβ to restore IPMK expression **(Fig. 6J)**. Moreover, verteporfin reduced FBXO22 association with pSMAD2/3 while enhancing FBXO22 binding to IPMK **(Fig. 6K)**. These results indicate that nuclear pSMAD2/3 displaces IPMK from the FBXO22-SCF complex.

Together, these findings identify TGFβ signaling as the upstream regulator of the IPMK-HDAC3 axis. TGFβ-induced nuclear pSMAD2/3 competitively engage FBXO22, thereby preventing FBXO22-CUL1-mediated ubiquitination and degradation of IPMK. Stabilization of IPMK preserves InsP₆ production, sustains HDAC3 activity, and mitigates ER stress by reducing CDK5RAP3.

### Short-term fasting in mice disrupts the TGF-β-IPMK-HDAC3 axis to induce ER stress

Our cell-based analyses defined a growth factor–dependent signaling hierarchy in which TGF-β stabilizes IPMK, sustains InsP₆-dependent HDAC3 activity, and restrains ER stress (**Fig. 7A**). We therefore asked whether in *in vivo* mouse intestine and enteroid model disruption of this pathway underlies fasting-induced ER stress.

Biochemical and immuno-histochemical analysis of mouse ileal epithelium revealed strong nuclear pSMAD2/3 staining under fed conditions, consistent with constitutive TGF-β signaling at steady state (**Fig. 7B-D**). The nuclear pSMAD2/3 signal was markedly reduced after a 12-h fast but restored following refeeding (**Fig. 7D**), indicating that fasting transiently suppresses TGF-β signaling *in vivo*.

Mouse intestinal enteroids basally activate TGF-β-mediated signaling. To determine whether loss of TGF-β signaling disrupts the IPMK–HDAC3 axis, we turned to mouse enteroids. Acute inhibition of TGF-β signaling with RepSox, a selective ALK5/TGF-β receptor inhibitor, rapidly reduced IPMK protein abundance (**Fig. 7E**) and caused a pronounced loss of HDAC3 enzymatic activity (**Fig. 7F**). Consistent with disruption of the IPMK–HDAC3 pathway, RepSox induced CDK5RAP3 expression, reduced RPL26 UFMylation, and robustly activated ER stress markers, including CHOP and IRE1 (**Fig. 7E**). These effects closely phenocopied the responses observed during growth factor deprivation in cell lines and fasting in mice, indicating that TGF-β signaling maintains epigenetic and proteostatic homeostasis in intestinal epithelium.

To test whether HDAC3 is the critical downstream effector, we next employed a chemical–genetic strategy. Acute PROTAC-mediated degradation of HDAC3 in otherwise untreated enteroids was sufficient to induce CDK5RAP3 expression, suppress RPL26 UFMylation, and activate ER stress markers (**Fig. 7G**), recapitulating the effects of TGF-β inhibition. Thus, HDAC3 activity loss represents a key nodal event linking growth factor signaling to ER stress.

Because IPMK degradation reduces cellular InsP₆ levels, we next asked whether restoration of InsP₆-dependent HDAC3 activity could rescue this phenotype. RepSox-treated enteroids were supplemented with cell-permeable InsP₆. Remarkably, InsP₆ suppressed CDK5RAP3 induction, rescued RPL26 UFMylation, and alleviated ER stress **(Supplementary Fig. 8A, B**). Notably, InsP₆ intervention did not restore pSMAD2/3 signaling **(Supplementary Fig. 8A, B**), indicating that stabilization of InsP₆-dependent HDAC3 activity is sufficient to bypass upstream TGF-β loss.

Finally, to genetically validate the pathway *in vivo*, we analyzed mice with intestinal epithelial cell–specific deletion of IPMK (IPMK*^IEC–/–^*). Under basal-fed conditions, control IPMK*^F/F^* mice exhibited intact RPL26 UFMylation and minimal expression of CDK5RAP3, CHOP, and IRE1, consistent with preserved ER homeostasis **(Fig 7H)**. In striking contrast, IPMK*^IEC–/–^* mice displayed constitutive induction of CDK5RAP3, elevated ER stress markers, and pronounced loss of RPL26 UFMylation even in the absence of fasting **(Fig 7H)**. Whereas a 12-hour fast robustly induced ER stress responses in IPMK*^F/F^* mice, fasting failed to further augment stress signaling in IPMK^IEC–/–^ animals, which remained locked in a maximal stress state **(Fig. 7I)**. Thus, genetic ablation of *IPMK* phenocopies fasting and renders IECs refractory to additional fasting-induced stress.

Together, these findings establish that fasting suppresses TGF-β signaling to destabilize the IPMK–HDAC3 axis, leading to InsP₆ depletion, HDAC3 inactivation, CDK5RAP3 induction, impaired ribosome-associated quality control, and ER stress.

## Discussion

Fasting engages a physiological quality-control mechanism that selectively eliminates genomically compromised epithelial cells. As illustrated in the final model **(Fig. 8)**, epithelial tissues, exemplified by the intestine, exist as mosaics of healthy and DNA-damaged cells. A brief fast converts metabolic stress into a selective pressure that eliminates the compromised population **(Fig. 1A–J)**, analogous to environmental stress–induced shedding of damaged leaves **(Fig. 8)**, while preserving overall tissue architecture. In the intestine, this process is executed through ER stress–dependent apoptosis **(Fig. 2)**, thereby reducing inflammatory burden **(Fig. 1L–O)**. Upon refeeding, regeneration proceeds from this purified cellular landscape, enabling healthy cells to repopulate the tissue. Thus, short-term fasting functions as a transient metabolic sieve that couples growth factor signaling to genomic integrity, ensuring that tissue renewal is seeded by the fittest cells.

**Figure 8.**
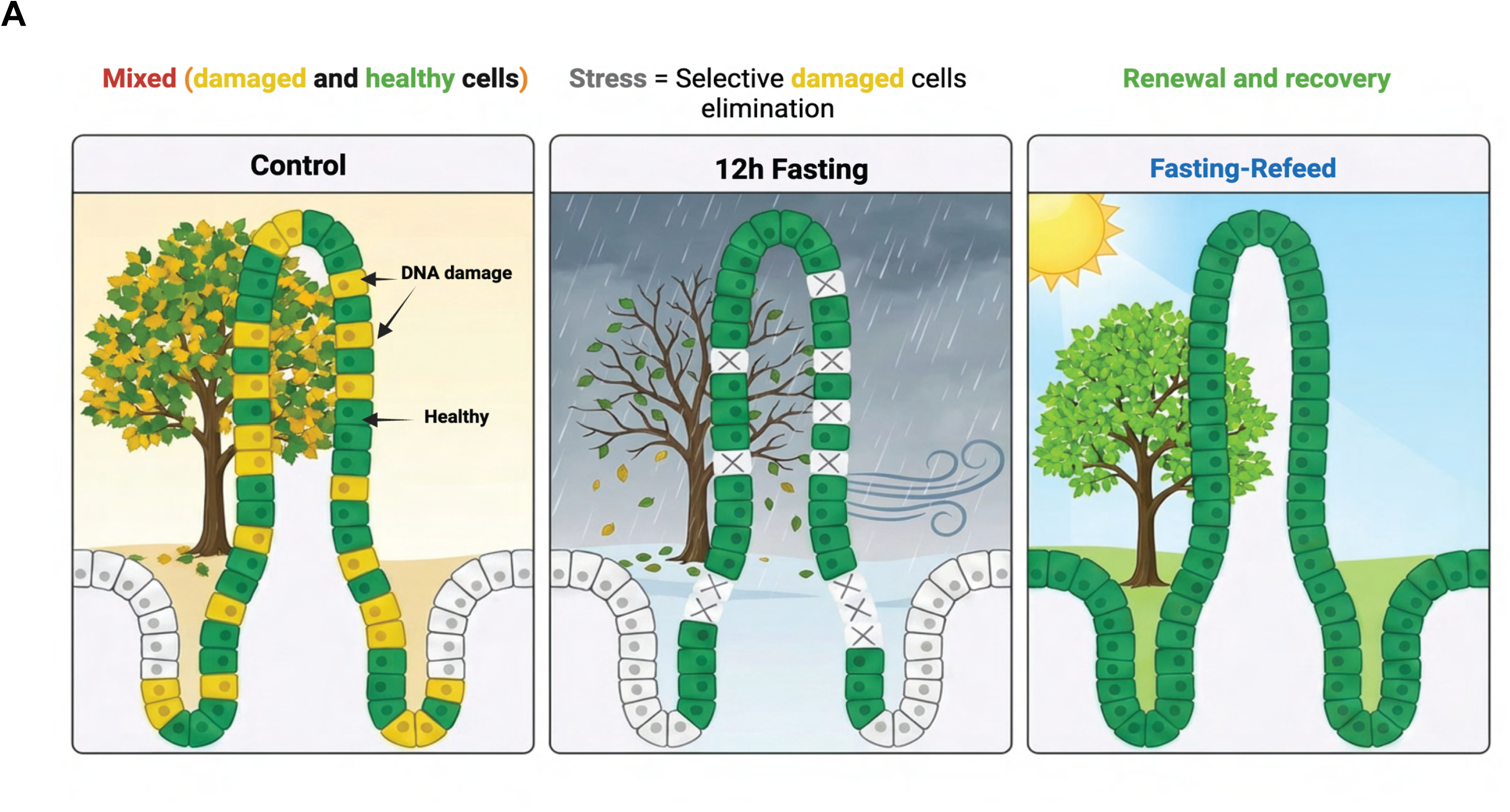
**(A)** Schematic representation of intestinal epithelium under control, 12 h fasting, and fasting–refeeding conditions. Under control conditions, the epithelium comprises a mosaic of healthy (green) and DNA-damaged (yellow) cells distributed along the crypt–villus axis. Following 12 h fasting, DNA-damaged cells are selectively eliminated (X) along the epithelium, while healthy cells are retained. Upon refeeding, epithelial architecture is restored through regeneration, resulting in a tissue predominantly composed of healthy cells. The background illustration depicts an analogous natural process observed in plants, in which stressed or damaged (yellow) leaves are shed under environmental stress (e.g., heavy wind), leaving behind healthy (green) leaves; subsequent favorable conditions promote regrowth and restoration of a healthy canopy. Schematic created in BioRender; Chatterjee, S., 2026; https://BioRender.com.

Mechanistically, we define a signaling cascade linking fasting-induced suppression of growth factor signaling to epigenetic and proteostatic remodeling **(Fig. 2-7)**. Fasting suppresses TGF-β signaling, enabling the FBXO22-Cullin1 ubiquitin ligase to target the inositol kinase IPMK for degradation **(Fig. 5A-H)**. Loss of IPMK depletes intracellular inositol hexaphosphate (InsP₆) **(Fig. 3J)**, an essential cofactor required for HDAC3 catalytic activation^10^. Consequently, HDAC3 inactivation triggers genome-wide increases in H3K27 acetylation and transcriptional induction of CDK5RAP3 **(Fig. 2G, H, 4D-G)**, while also promoting acetylation-dependent stabilization of the CDK5RAP3 protein **(Fig. 2K)**. Elevated CDK5RAP3 disrupts ribosome-associated UFMylation of RPL26, leading to impaired ER proteostasis and activation of ER stress **(Supplementary Fig. 2B**).

Importantly, ER stress alone does not indiscriminately induce apoptosis. Instead, it acts synergistically with pre-existing genomic instability to push DNA-damaged cells beyond an apoptotic threshold **(Supplementary Fig. 1G, 2D**). Cells lacking this genotoxic priming remain largely resistant to ER stress-induced death. This selective vulnerability provides a mechanism by which fasting removes compromised cells while preserving tissue function.

Our findings further reveal an unexpected link between growth-factor signaling and epigenetic control of proteostasis. Under nutrient-replete conditions, constitutive TGF-β signaling stabilizes IPMK by redirecting the FBXO22–Cullin1 ubiquitin ligase toward phosphorylated SMAD2/3 substrates in the nucleus **(Fig. 6A-K)**. Fasting disrupts this balance, allowing FBXO22 to engage chromatin-associated IPMK and promote its degradation. Because IPMK-derived InsP₆ serves as an essential cofactor for HDAC3 catalytic activation^10^, loss of IPMK during fasting depletes cellular InsP₆ and inactivates HDAC3. In this manner, loss of growth-factor signaling is converted into a metabolic checkpoint that suppresses HDAC3 activity and initiates downstream stress responses.

A notable feature of this pathway is the central role of higher-order inositol phosphate metabolism in regulating epigenetic signaling. Our previous work established that IPMK associates with nuclear HDAC3 to regulate histone deacetylation^10,23^, whereas the present study shows that depletion of IPMK-derived InsP₆ during fasting broadly impairs HDAC3 activity toward both histone and non-histone substrates. Because IPMK is predominantly nuclear, its degradation collapses the nuclear InsP₆ pool that serves as a major intracellular reservoir for HDAC3 activation. One possibility is that HDAC3 acquires InsP₆ in the nucleus, where cofactor binding licenses catalytic activity prior to redistribution to cytoplasmic compartments. Alternatively, nuclear InsP₆ may diffuse to extra nuclear sites to activate local HDAC3 pools associated with non-histone substrates such as CDK5RAP3. Under either model, degradation of nuclear IPMK would limit InsP₆ availability and broadly restrict HDAC3 activity across cellular compartments.

This defines a pathway in which fasting suppresses growth factor signaling to drive IPMK loss, InsP₆ depletion, HDAC3 inactivation, CDK5RAP3 stabilization, impaired UFMylation, and ER stress.

Moreover, this IPMK–InsP₆–HDAC3–CDK5RAP3-ER stress signaling axis engaged during fasting appears conserved across multiple organs **(Supplementary Fig. 2E**), suggesting that this pathway represents a general metabolic mechanism of tissue quality control.

Because the accumulation of DNA-damaged cells is a hallmark of aging tissues and fuels chronic inflammation and tumor initiation, coupling nutrient availability to the selective removal of genomically unstable cells may represent an adaptive strategy to preserve tissue integrity. Using IBD as a model of chronic inflammation, we show that enhancing epithelial quality control through short-term fasting reduces inflammation level in DSS colitis model **(Fig 1L-O, Supplementary Figure H)**. Fasting may therefore transiently activate a growth factor-sensitive damage-surveillance program that limits the persistence of DNA-damaged cells. Such mechanisms provide a unifying framework for the beneficial effects of fasting on tissue homeostasis, inflammation, and age-associated disease, like cancer, and suggest that metabolic state can act as a physiological regulator of tissue composition by selectively purging damaged cells to preserve long-term organ integrity.

## Methods

### Cell culture

Human colon tumor (HCT116), mouse embryonic fibroblast (MEF), A549, HEK293, B16F10, MDA-MB-231, HepG2, MCF7, and CaCo-2 cell lines were cultured in McCoy’s 5A medium, Dulbecco’s Modified Eagle’s Medium (DMEM), RPMI, or F12/K medium, as appropriate for each cell type. DMEM (Cat. No. 11965092 Gibco), F12/K (Cat. No. 21127-022), and RPMI (Cat. No. 12633012) media were supplemented with 10% fetal bovine serum (FBS) (Gibco, Cat. No. A52567-01), 2 mM L-glutamine, 100 U/mL penicillin, and 100 µg/mL streptomycin and were used for culturing MEF and A549 cells. McCoy’s 5A medium (Cytiva Cat. no. SH30200.02), used for HCT116 cells, was supplemented with 10% FBS and 100 U/mL penicillin-streptomycin.

*IPMK*, *HDAC1, and HDAC3* knockout HCT116 cells were generated using CRISPR/Cas9 gene-editing technology. Following transfection, clonal populations were isolated, expanded, and screened for IPMK expression^10^.

### Treatment of cells with various physiological stressors

To induce stress related to heat shock, 1 × 10⁶ HCT116 cells were seeded into a six-well plate. The following day, the cells were exposed to 42°C for 60 minutes using a pre-warmed, sterilized water bath^32^.

To induce stress related to cold shock, 1 × 10⁶ HCT116 cells were seeded into a six-well plate. The following day, the cells were exposed to 4°C for 60 minutes^33^.

To induce osmotic stress, 1 × 10⁶ HCT116 cells were seeded into a six-well plate. On the following day, the cells were treated with regular culture media supplemented with 500 mM D-sorbitol for 60 minutes^34^.

To induce oxidative stress, 1 × 10⁶ HCT116 cells were seeded into a six-well plate. On the following day, the cells were treated with regular culture media supplemented with 500 μM H_2_O_2_ for 3 hr^35^.

To induce UV-induced DNA damage-related stress, 1 × 10⁶ HCT116 cells were seeded into a six-well plate. The following day, the cells were exposed to UV irradiation at a dose of 70 J/m² for 60 minutes in regular culture media^36^.

To induce amino acid (AA) deprivation-related nutritional stress, 1 × 10⁶ HCT116 cells were seeded into a six-well plate. On the following day, the cells were incubated in Hanks’ Balanced Salt Solution (HBSS) supplemented with 10% FBS for 3 hours.

To induce growth factor deprivation-related nutritional stress, 1 × 10⁶ HCT116 cells were seeded into a six-well plate. The following day, the cells were incubated in McCoy’s/DMEM medium without FBS supplementation, as FBS serves as the primary source of growth factors in standard cell culture conditions.

### Generation of IPMK antibody

A custom anti-IPMK antibody was generated in collaboration with Pro-Sci. The anti-rabbit IPMK antibody had been raised against human IPMK amino acids 295–311 (SKAYSRHRKLYAKKHQS). These sequences are also closely conserved in mice **SK**M**Y**A**RHRK**I**Y**T**KKH**H**S,** hence react to both mouse and human IPMK^10^.

### Western Blot analysis

Cell lysates were prepared through syringe flush by using RIPA Lysis buffer (50 mM Tris pH 7.5, 150mM NaCl, 1% NP-40, 0.1% SDS, 0.5% sodium deoxycholate, 1mM EDTA plus protease and phosphatase inhibitor). Samples were centrifuged at 15,000g for 10 min, and the protein concentration of the supernatant was measured. Proteins were resolved by SDS-polyacrylamide gel electrophoresis (NuPAGE Bis-Tris Midi GEL, Life Technologies, Cat #: WG1402BX10) and transferred to Immobilion-P PVDF (Millipore-Sigma, Cat #: IPVH00010) transfer membranes. The membranes were incubated with primary antibody diluted in 3% BSA in Tris-Buffered Saline with Tween 20 (20 mM Tris-HCl, pH 7.4, 150 mM NaCl, and 0.02% Tween 20) overnight incubation at 4°C. Respective antibodies for western blot anti-HDAC3 (Santa Cruz Biotechnology, Cat #: sc-376957), IPMK (Generated in our lab), anti-β actin (ProteinTech, Cat #: 81115-1-RR), anti-lamin (ProteinTech, Cat #: 12987-1-AP), anti-NCoR2 (Bethyl Laboratories, Cat #: A301-148A), anti-Ncor1 (Cell Signaling Technology, Cat #: 5948), anti-TBL2 (ProteinTech, Cat #: 26686-1-AP), Anti-SMRT (ProteinTech, Cat #: 14638-1-AP), anti-GPS2 (Bethyl Laboratories, Cat #: A304-048A), anti-H3 (Cell Signaling Technology, Cat #: 12648), anti-Pan Ac (Cell Signaling Technology, Cat #: 9441), anti-H3k9Ac (Cell Signaling Technology, Cat #: 9649), anti-H3k27Ac (Abcam, Cat #: ab4729), anti-H4k16Ac (Cell signaling Technology, Cat #: 13534S), anti-GST, anti-Flag (ProteinTech, Cat #: 20543-1-AP), anti-myc (Invitrogen, Cat #: 46-0709) anti-CDK5RAP3 (ProteinTech, Cat #: 11007-1-AP),anti-CDK5RAP3 (Bethyl, Cat no. A300-870), anti-CHOP (ProteinTech, Cat #: 66741-1-IG), anti-IRE1 (ProteinTech, Cat #: 27528-1-AP),anti-Calereticulin (Cell signaling Technology, Cat #: 12238S), Cleaved Caspase 3 (Cell signaling Technology, Cat #: 9664S), anti-∀H2X (Abcam, Cat# Ab11174), anti-Myc (Invitrogen, Cat #:46-0709), anti-LC3B (Cell Signaling Technology, Cat #: 3868T), anti-ChK2 (Cell signaling Technology, Cat #:2662S), anti-pChK2 (Cell signaling Technology, Cat #:2197T), anti-IL-1β (Cell signaling Technology, Cat #:12426T), anti-EpCAM (Invitrogen, Cat #:13-5791-82), anti-FBX022 (ProteinTech, Cat #: 13606-1-AP), anti-Cul1(ProteinTech, Cat #:82474-1-RR), anti-FBX02(ProteinTech, Cat #: 14590-1-AP, anti-His(ProteinTech, Cat #:66005-1-Ig), anti-SKP1(ProteinTech, Cat #: 10990-2-AP), anti-Pan Ubiquitination (ProteinTech, Cat #:10201-2-AP) and anti-K48 ubiquitin (Cell signaling Technology, Cat #:8081T). Following primary antibody incubation, the PVDF membrane was washed three times with TRIS-buffered saline/Tween-20 and incubated with the respective HRP-conjugated secondary antibody (ECL, Cat #s: NA934V and NA931V). The bands were visualized by chemiluminescence (Super Signal West Pico, Pierce, Cat #: 34579). The depicted blots are representative replicates selected from at least three experiments. Densitometric analysis was performed using ImageJ software. Blots for acetylated histones were stripped using Restore Western Blot stripping buffer (Thermo Scientific, Cat # 21059) then reprobed with H3 or H4 antibody (Cell Signaling Technology, Cat #: 12648S, 2935S) at room temperature for 2 hours, followed by secondary antibody incubation and development^10^.

### SILAC labeling and mass spectrometry

For SILAC labeling, HCT116 cells were cultured in suspension for eight doublings. One population was maintained in ‘light’ complete McCoy’s 5A medium, while the other was grown in ‘heavy’ SILAC McCoy’s 5A medium (Thermo Scientific, Cat. No. 88441), in which L-arginine was replaced with R6-labeled L-arginine (Cambridge Isotope Laboratories, Cat. No. CNLM-90007-CA). Samples were digested overnight with trypsin (Pierce) following reduction and alkylation with DTT and iodoacetamide (Sigma–Aldrich). The samples then underwent solid-phase extraction cleanup with an Oasis HLB plate (Waters) and the resulting samples were injected onto a QExactive HF mass spectrometer coupled to an Ultimate 3000 RSLC-Nano liquid chromatography system. Samples were injected onto a 75 μm i.d., 15-cm long EasySpray column (Thermo) and eluted with a gradient from 0-28% buffer B over 90 min with a flow rate of 250 nL/min. Buffer A contained 2% (v/v) ACN and 0.1% formic acid in water, and buffer B contained 80% (v/v) ACN, 10% (v/v) trifluoroethanol, and 0.1% formic acid in water. The mass spectrometer operated in positive ion mode with a source voltage of 2.5 kV and an ion transfer tube temperature of 275 °C. MS scans were acquired at 120,000 resolution in the Orbitrap and up to 20 MS/MS spectra were obtained for each full spectrum acquired using higher-energy collisional dissociation (HCD) for ions with charges 2-8. Dynamic exclusion was set for 20 s after an ion was selected for fragmentation.

Raw MS data files were analyzed using Proteome Discoverer v3.0 (Thermo), with peptide identification performed using Sequest HT searching against the human protein database from UniProt. Fragment and precursor tolerances of 10 ppm and 0.6 Da were specified, and three missed cleavages were allowed. Loss of N-terminal methionine and N-terminal acetylation were set as protein variable modifications. Carbamidomethylation of Cys was set as a fixed peptide modification, with oxidation of Met, methylation of Lys and Arg, dimethylation of Lys and Arg, trimethylation of Lys, acetylation of Lys, and propionylation of Lys set as peptide variable modifications. The false-discovery rate (FDR) cutoff was 1% for all peptides.

### HDAC (histone deacetylase) activity

To study global HDAC activity, we used an HDAC activity assay kit (Active Motif, Cat #: 56200). In brief, we lysed cells in a low-salt lysis buffer (50 mM Tris pH 7.5, 50 mM potassium acetate, 5 % v/v glycerol, 0.3 % v/v Triton X-100, one tablet of Roche complete protease inhibitor), followed by centrifugation at 15,000 g to isolate the supernatant. About 20 µg μg of protein from UT and GFD HCT116 cell lysates were incubated with the substrate provided with the kit, and then fluorescence was measured using a plate reader. To analyze deacetylase activity of specific Class I HDAC members, HDAC1, 2, 3, and 8 were individually immuno-purified, followed by an activity assay using the following kits: HDAC Assay Kit (Active Motif, Cat#: 56200 for HDAC1, 2, and 8) and the HDAC3 Assay Kit (BPS Bioscience, Cat#: 10186-628). Since high salt concentration dissociates higher-order inositol phosphates (HOIPs) from HDAC1/3, which are crucial for their deacetylase activity, cells were lysed in salt-free (NaCl-free) lysis buffer (50 mM Tris pH 7.5, 50 mM potassium acetate, 5% v/v glycerol, 0.3% v/v Triton X-100, and one tablet of Roche complete protease inhibitor). Then, the lysate was centrifuged at 15,000 g for 10 min, and the protein concentration of the supernatant was measured. Equal amounts, around 20 µg total protein from the lysate, were incubated against anti-HDAC1 (Santa Cruz Biotechnology, Cat # sc-81598), anti-HDAC2 (Santa Cruz Biotechnology, Cat #: sc-7899), anti-HDAC3 (Santa Cruz Biotechnology, Cat #: sc-81600) and anti-HDAC8 (Biolegend, Cat #: 685504) antibodies, respectively, at 4°C for 1 hour followed by capturing antibody with EZview A/G beads (Millipore Sigma, Cat #: E3403). Beads were washed 3x with an ice-cold reaction buffer (supplied with the kit). After a specific substrate reaction, fluorescence was measured using a Spectra max iD5 plate reader (Molecular Devices). IgG was immunoprecipitated from Wild Type HCT116/MEF or HEK293 cells (WT) and used as a negative control. In each experiment, separate IgG immunoprecipitation was performed, and the data was expressed as a fold change relative to the respective IgG control. Although slight variations in IgG values were observed across experiments; the overall data range differed slightly in each case but consistently showed a similar trend. All measurements were performed in triplicate, and data was analyzed using GraphPad Prism (version 6.0, GraphPad Software, Inc)^10,23,37^.

### HAT (histone acetyltransferase activity)

HAT activity was measured by following the HAT Assay Kit (Abcam, Cat #: ab-204709) protocol. Cells were lysed in salt-free (NaCl-free) lysis buffer (50 mM Tris pH 7.5, 50 mM potassium acetate, 5% v/v glycerol, 0.3 % v/v Triton X-100, Roche complete protease inhibitor). Samples were centrifuged at 15,000 g for 10 min, and the protein concentration of the supernatant was measured. Equal amounts of (around 20 µg of total protein from the lysate) were incubated against anti-p300 antibody (Cell Signaling Technology, Cat #: 86377) at 4°C for 90 min, followed by capturing the antibody with EZview A/G beads (Millipore-Sigma, Cat #: E3403). Beads were washed 3x with an ice-cold reaction buffer (supplied with kit). Immunoprecipitated P300 was used as a protein source. Absorbance was measured after specific substrate reaction by using a Spectramax iD5 plate reader (Molecular Devices). IgG was immunoprecipitated from wild-type HCT 166 cells (WT) and used as a negative control. In each experiment, separate IgG immunoprecipitation was performed, and the data was expressed as a fold change relative to the respective IgG control. Although slight variations in IgG values were observed across experiments, the overall data range differed slightly in each case but consistently showed a similar trend. All measurements were performed in triplicate, and data were analyzed using GraphPad Prism (version 6.0, GraphPad Software, Inc)^10,23^.

### InsP_6_ rescues HDAC3 activity

Cells were lysed in salt-free (NaCl-free) lysis buffer (50 mM Tris pH 7.5, 50 mM potassium acetate, 5% v/v glycerol, 0.3 % v/v Triton X-100, and one Roche complete protease inhibitor tablet). Samples were centrifuged at 15,000 g for 10 min, and the protein concentration of the supernatant was measured. Equal amounts, 20 µg total protein from the lysate, were incubated against anti-HDAC3 antibodies, respectively, at 4°C for 1 hr, followed by capturing antibodies with EZview A/G beads (Millipore Sigma, Cat #: E3403).

Beads were washed 3x with an ice-cold reaction buffer (supplied with the kit). Following washing, beads were resuspended in reaction buffer, and then beads were incubated with 0.01 µM, 0.1 µM, 1 µM, 5 µM, and 10 µM concentrations of InsP_6_ (Sigma Aldrich Cat #: P8810-100G) for 1 hr at room temperature. After incubation, beads were washed 3x with a reaction buffer. HDAC3 activity was measured by the addition of a fluorogenic substrate provided by the HDAC3 Assay Kit (BPS, Cat #: 10186-628) and read using a Spectramax iD5 plate reader (Molecular Devices). IgG was immunoprecipitated from wild-type untreated cells (WT) and used as a negative control. In each experiment, separate IgG immunoprecipitations were performed, and the data were expressed as fold change relative to the respective IgG control. Although slight variations in IgG values were observed across experiments, the overall data range differed slightly in each case but consistently showed a similar trend. All measurements were performed in triplicate data and analyzed using GraphPad Prism (version 6.0, GraphPad Software, Inc)^10,23,37^.

### InsP_6_ rescues HDAC3 activity and H3K27ac in the cell-free nucleus

Approximately 5 × 10⁶ cells were washed twice with ice-cold PBS and resuspended in swelling buffer (10 mM Tris–HCl, pH 7.5; 2 mM MgCl₂; 3 mM CaCl₂) supplemented with protease and phosphatase inhibitors. Cells were allowed to swell on ice for 10 min, followed by washing with swelling buffer containing 10% glycerol. Nuclei were isolated by lysing cells in nuclear lysis buffer (10 mM Tris–HCl, pH 7.5; 2 mM MgCl₂; 3 mM CaCl₂; 10% glycerol; 1% Igepal NP-40; protease and phosphatase inhibitors) for 15 min on ice. The nuclear pellet was subsequently washed twice with the same buffer.

Isolated nuclei were permeabilized with 5% digitonin in lysis buffer for 15 min, and then after two times washing in lysis buffer and one washing with reaction buffer, incubated with 1 μM inositol hexakisphosphate (InsP₆) for 60 min at room temperature in the reaction buffer provided with the HDAC3 activity assay kit (BPS Bioscience, Cat. #10186-628).

Following incubation, nuclei were lysed in salt-free lysis buffer (50 mM Tris–HCl, pH 7.5; 50 mM potassium acetate; 5% glycerol; 0.3% Triton X-100; supplemented with one Roche Complete protease inhibitor tablet). Lysates were centrifuged at 15,000 × g for 10 min, and the protein concentration of the supernatant was determined by running a western blot for evaluation of H3K27ac. Equal amounts of protein (20 μg) were subjected to immunoprecipitation with anti-HDAC3 antibodies at 4 °C for 1 h, followed by capture with EZview Protein A/G agarose beads (Millipore Sigma, Cat. #E3403). The beads were washed three times with ice-cold reaction buffer supplied with the assay kit.

HDAC3 enzymatic activity was measured by adding the fluorogenic substrate provided in the HDAC activity assay kit (BPS Bioscience, Cat. #10186-628) and fluorescence was quantified using a SpectraMax iD5 plate reader (Molecular Devices) according to the manufacturer’s instructions.

For negative controls, IgG immunoprecipitation from untreated wild-type cells was performed in parallel in each experiment. HDAC3 activity was expressed as fold change relative to the corresponding IgG control. Although minor variations in IgG baseline values were observed between experiments, the relative trends remained consistent across replicates.

All experiments were performed in biological triplicates, and data were analyzed using GraphPad Prism (version 6.0).

### Plasmids and transient transfection

pMX-myc, pMX-*IPMK-WT-*myc, pMX-*IPMK-KD*-myc, cmv-FLAG, cmv-DAD-V5-FLAG, cmv-*CDK5RAP3*-GFP (Sino Biological. Cat no. MG-52956-ANG), pcmv3-GFP-Control (Sino Biological. Cat no. CV026), pcmv-CDK5RAP3-myc (Sino Biological. Cat no. MG-52956-NM), and cmv-myc plasmids were transiently transfected into HCT116 cells using Lipofectamine 3000 (Cat. No. L3000001, Thermo Scientific), according to the manufacturer’s protocol.

### Lentivirus Transfection and Stable Cell Line Development

A total of 0.5 × 10⁶ HDAC3 cells were transfected with GFP-tagged lentiviral constructs, either^Lenti^-HDAC3^WT^ (CMV>Myc/V5/3xGS/hHDAC3[NM_003883.4]:WPRE:EF1A>EGFP) or^Lenti^-HDAC3^HOIP^(pLV[Exp]-CMV>Myc/V5/3xGS/hHDAC3[NM_003883.4](R265P)(L266M):WPRE:EF1A>EGFP) using Polybrene as the transfection reagent. Following 48 hours of incubation.

### Inhibitors

For the inhibition of TGF-β signaling, cells were treated with 1 μM Vactosertib (Cat. No. 19928 MCE chemicals) or 50 μM RepSoX (Cat no. HY-13012 MCE Chemical) overnight (approximately 16 hours) in complete growth media supplemented with serum.For the enteroid system, RepSOX treatment was administered for 6 hr at a dose of 50 μM. Longer treatment durations were not used because enteroids exhibited significant loss of viability or morphological changes^38^.

To inhibit proteasomal degradation, both untreated control cells and GFD-treated cells were exposed to 10 μM MG132 (Cat. No. HY-13259 MCE chemicals) for 3 hours under standard culture conditions.

For the inhibition of histone acetyltransferase P300, cells were treated with 10 μM A845 (Cat. No. HY-107455 MCE chemicals) and incubated overnight (12 hr) in serum-free GFD media, ensuring the selective inhibition of P300 activity in nutrient-deprived conditions.

To inhibit Tip60/KAT5 acetyltransferase activity, cells were treated with 35 μM MG149 (Cat. No. HY-15887 MCE chemicals) for overnight incubation (12hr) in serum-free GFD media to allow for sufficient pharmacological inhibition under nutrient stress conditions.

Wild-type (WT) cells were treated with 2 mM Suberoylanilide Hydroxamic Acid (SAHA), a broad-spectrum histone deacetylase (HDAC) inhibitor obtained from Sigma-Aldrich (Cat. No. SML0061-5MG), and incubated overnight in complete media to assess the effects of HDAC inhibition on cellular processes^23^.

For autophagy inhibition, cells were treated with 100 nM Bafilomycin A (Cayman Chemicals, Cat. No. 11038) for 3 hours. This treatment was performed under standard culture conditions to block autophagosome-lysosome fusion and inhibit lysosomal acidification^35,39^.

To inhibit proteasomal degradation, both untreated control cells and GFD-treated cells were exposed to 1 μM Bortisimib (Cat. No. HY-10227, MCE chemicals) for 3 hours under standard culture conditions.

To inhibit proteasomal degradation, both untreated control cells and GFD-treated cells were exposed to 10 μM TAK243 (Cat. No. HY-100487, MCE chemicals) for 3 hours under standard culture conditions.

To inhibit proteasomal degradation, both untreated control cells and GFD-treated cells were exposed to 1 μM Pevanostat (Cat. No. HY-10484, MCE chemicals) for 3 hours under standard culture conditions.

For the enteroid system, 1 μM Pevanostat was administered along with RepSOX for 6 hr. For ULK1/2 inhibition, cells were treated with 10 μM SBI-0206965 (Cayman Chemicals, Cat. No. 18477) for 24 hours. This treatment was performed under standard culture conditions.

### siRNA transfection

HCT116 cells were seeded into six well plates at a density of 30,000 cells per well one day prior to siRNA transfection. For each gene, a siRNA pool was employed, comprised of equimolar amounts of four individual ON-TARGETplus siRNA duplexes (Dharmacon) targeting distinct coding regions of the targeted mRNA. A negative control siRNA (siScramble) was employed in all silencing experiments. Cells were transfected using RNAiMAX (ThermoFisher Scientific) according to the manufacturer’s instructions. Briefly, for each well to be transfected, 10 pmol siRNA pool was combined with 3 uL RNAiMAX in 500 μL serum-free medium. The mixture was incubated for 20 min at room temperature before being dispensed into the cell culture plate. Cell culture medium was replaced with complete medium on the subsequent day. Three days post-transfection, cells were subjected to complete GFD or 2 μM Vactosertib treatment for a duration of 12 h prior to protein harvest. Target gene silencing was confirmed by Western blot using the antibodies.

### HDAC3 PROTAC treatment

The HDAC3 PROTAC used in this study was originally developed and characterized in the laboratory of Dr. Frank J. Dekker in the Netherlands and is denoted **P7** in the prior publication. For all experimental purposes, the HDAC3 PROTAC compound was initially dissolved in DMSO (Dimethyl Sulfoxide) to prepare a stock solution, ensuring optimal solubility and stability^40^.

Based on prior observations indicating that higher concentrations of this PROTAC exhibit off-target effects, particularly on HDAC8 protein levels, a carefully optimized lower concentration was selected to ensure specificity toward HDAC3. Accordingly, wild-type (WT) HCT116 cells were treated with the HDAC3 PROTAC at a final concentration of 41.2 nM.

The treatment was performed for various time points to study the time-dependent effects of HDAC3 degradation. Cells were exposed to the PROTAC for 4, 8, 12, 18, and 24 hours under standard cell culture conditions.

To further investigate the reversibility and recovery dynamics following PROTAC treatment, a washout experiment was performed after 24 hours of continuous exposure to the compound. The PROTAC-containing media was removed, cells were washed thoroughly with fresh media, and then incubated in PROTAC-free media for an additional 2, 4, and 6 hours, respectively.

For the enteroid culture system, HDAC3 PROTAC was administered at a final concentration of 41.2 nM for 6 hr. Longer treatment durations were not used because enteroids exhibited significant loss of viability and morphological changes. This experimental design allowed for the assessment of both the efficacy of HDAC3 degradation and the potential recovery of protein levels upon PROTAC removal.

### Growth factor treatment (IGF, TGF, VEGF, FGF, EGF, PDGF)

Wild-type (WT) HCT116 cells were initially cultured in serum-free, growth factor-deprived (GFD) media for 12 hours (overnight). After the overnight deprivation, cells were treated with a panel of recombinant human growth factors, including IGF (26 nM), VEGF (5 nM), EGF (5 nM), FGF (5 nM), PDGF (5 nM), and TGF-β (200 pM), for 3 hours under standard culture conditions. The growth factors were prepared following the manufacturer’s instructions to ensure their biological activity. After the 3-hour treatment, cells were lysed using lysis buffer, and the resulting protein extracts were collected for downstream biochemical analyses.

### Immunoprecipitation

To analyze endogenous binding, HCT116 cells were lysed in salt-free lysis buffer (50 mM Tris pH 7.5, 50 mM potassium acetate, 5% v/v glycerol, 0.3 % v/v Triton X-100, and one Roche complete protease inhibitor tablet). For the endogenous immunoprecipitation (IP), respective antibodies against HDAC3 were used, followed by Western blot of the binding partners. In brief, the IP was performed from 500 μg of protein lysate. Protein lysates were incubated for 2 hr at 4°C with respective antibodies, then the antibody-protein complex was captured with EZview A/G beads (Millipore Sigma, Cat #: E3403). Beads were pelleted and washed with lysis buffer 3x, followed by elution in sample buffer. Immunoprecipitated samples were resolved on a NuPAGE Bis-Tris gel, followed by western blotting^10^.

### ChIP-qPCR

Approximately 7×10^6^ cells were fixed with 1% final volume from fresh 16% stock formaldehyde (Sigma-Aldrich, Cat#: F8775) at room temperature for 10 min followed by ChIP using iDEAL ChIP assay Kit from Diagenode (Cat# C01010051). Cells were then harvested and lysed in 500 mL of ChIP lysis Buffer (50 mM Tris-HCl pH 8.0, 5 mM EDTA, 150 mM NaCl, 0.5% Triton X-100, 0.5% SDS, 0.5% NP-40, 1 mM sodium Technology, Cat#13534), HDAC3 (Santa Cruz Biotechnology, Cat# 39717) and HDAC1 (Santa Cruz Biotechnology, Cat# 81598X) 4C overnight. Then, the lysate was incubated with Protein G magnetic bead (provided in kit) for 1 h at 4C. The beads were washed sequentially with a wash buffer provided in the kit. The immune complexes were eluted with 75 mL of elution buffer (1% SDS, 0.1 M NaHCO3) twice at 65C for 30 min. After elution, the cross-link was reversed by adding NaCl and incubated together with Proteinase K provided with the kit overnight at 65C. ChIP DNA was purified using the ChIP DNA purification kit provided with the kit. The purified DNA was analyzed on a StepOnePlus using the power SYBR Green Master Mix. The results are presented as a fold change in respect with IgG after calculating the percentage of input. qPCR analyses were done in triplicate. We used CDK5RAP3 5′-GAGCGTAAACCCTGATTGGC-3′ and 5′-GCCCTACCACCAATCCAGT-3′ for human and CDK5RAP3 5′-GAGTGTTGCTTCCTCTCCCT-3′ and 5′-GTCACAGGTCTCTTGAGGCT-3′ for mice^10^.

### CUT&RUN sample preparation

CUT&RUN assays were performed using a modified protocol as follows. Briefly, approximately 1 × 10ª cells were harvested, washed twice with ice-cold PBS, and crosslinked with formaldehyde and DSG, followed by quenching with 1 M glycine. Cells were washed twice with wash buffer (20 mM HEPES pH 7.5, 150 mM NaCl, 0.5 mM spermidine, supplemented with protease inhibitors) and resuspended in wash buffer. Concanavalin A magnetic beads were activated prior to use by washing twice with bead activation buffer (20 mM HEPES pH 7.5, 10 mM KCl, 1 mM CaCl2, 1 mM MnCl2) and kept on ice. Cells were then incubated with activated beads for 20 min at room temperature to immobilize nuclei, followed by permeabilization using cell permeabilization buffer consisting of wash buffer supplemented with 0.01% digitonin. Bead-bound nuclei were incubated overnight at 4 °C with the indicated antibody in antibody incubation buffer composed of permeabilization buffer supplemented with 0.5 mM EDTA. After antibody binding, samples were washed and incubated with pAG-MNase for targeted chromatin cleavage. Digestion was initiated by adding CaCl2 incubation buffer (CAB) containing 3.5 mM HEPES pH 7.5, 10 mM CaCl2, and 0.05% digitonin, and samples were incubated at 4 °C to allow controlled MNase activation. The reaction was terminated by addition of stop buffer (170 mM NaCl, 20 mM EGTA, 0.05% digitonin, RNase A, and glycogen), and released DNA fragments were collected from the supernatant. Crosslinks were reversed by incubation with SDS and Proteinase K at 55 °C overnight, followed by purification using a column-based DNA cleanup method. Purified DNA was quantified using a Qubit fluorometer and stored at –20 °C prior to library preparation and sequencing.

Eluted DNA from CUT&RUN was used for library construction with the CUTANA™ CUT&RUN Library Prep Kit (EpiCypher, Cat. #14-1001 and #14-1002), according to the manufacturer’s instructions.

### Analysis of CUT&RUN data

Sequencing reads were trimmed to remove adapter sequences and reads shorter than 20 bp. Reads were then aligned to either the human (Ensembl GRCh38) or mouse (Ensembl GRCm39) reference genome with bowtie2 (v 2.4.4; Langmead and Salzberg, 2012). For spike-in normalization, reads were also aligned tothe *Escherichia coli* K12 MG1655 reference genome (Ensembl Bacteria ASM584v2). Aligned reads were subsequently filteredto remove reads overlapping ENCODE blacklisted regions. Peaks were called on filtered alignments using MACS2 (v 2.2.7.1; Gaspar, 2018). Normalized coverage tracks were generated using the genomecov function from bedtools (v 2.30.0; Quinlan and Hall, 2010) and converted to bigWig format using bedGraphToBigWig (v 2.10; Kent et al., 2010). Signal distributions across genes were calculated using the computeMatrix module from deepTools (v 3.5.1; Ramirez et al., 2016) and visualized with the plotHeatmap function. To determine differential peaks, we used the R packages DiffBind (v 3.16.0; Stark and Brown, 2011) and DESeq2 (v 1.46; Love et al., 2014) with threshold values of absolute log2 fold change greater than 1.0 and FDR < 0.05 between group comparisons. The differential peaks were then annotated with nearby genes with the R package ChIPseeker (v 1.42.1; Yu et al., 2015). Annotated peaks were plotted on normalized coverage tracks using pyGenomeTracks (v 3.9; Lopez-Delisle et al., 2021)^41–45^.

### Cell Lysis and Fractionations

Approximately 7×10^6^ cells were harvested and washed three times with ice-cold phosphate-buffered saline (PBS) to remove residual medium. The cell pellet was lysed in ice-cold cytoplasmic lysis buffer containing 50 mM HEPES-KOH (pH 7.5), 140 mM NaCl, 1 mM EDTA, 10% (v/v) glycerol, 0.5% (v/v) NP-40, 0.25% (v/v) Triton X-100, and 1 mM DTT, supplemented with a 1× protease inhibitor cocktail (Roche). The lysate was incubated on ice for 10 minutes and centrifuged at 1100 × g for 5 minutes at 4 °C. The resulting supernatant was collected as the cytoplasmic fraction, while the pellet, containing crude nuclei, was retained for further processing.

The nuclear pellet was washed three times with cytoplasmic lysis buffer to eliminate cytoplasmic contamination. Subsequently, the pellet was resuspended in nuclear lysis buffer composed of 10 mM Tris-HCl (pH 7.5), 200 mM NaCl, 1 mM EDTA, and 0.5 mM EGTA, supplemented with 1× protease inhibitor cocktail. The nuclei were gently mixed by pipetting up and down ten times and incubated on ice for 10 minutes. Following incubation, the suspension was centrifuged at 1100 × g for 10 minutes at 4 °C. The supernatant was collected as the nuclear fraction, and the remaining pellet, containing crude chromatin, was washed three times with nuclear lysis buffer. The resulting unsheared chromatin fraction was then used for downstream applications ^10^.

### Chromatin Immunoprecipitation and Mass Spectrometry

For immunoprecipitation of chromatin-associated IPMK, cells were transfected to overexpress myc-tagged IPMK plasmid. Following drug treatment with growth factor deprivation (GFD) and MG132, chromatin was isolated using the protocol described above. The chromatin fraction was subsequently subjected to immunoprecipitation using anti-Myc Trap agarose beads (Cat. #YTA20; Chromotek). After three times washing with lysis buffer to remove nonspecific interactions, bead-bound complexes were processed for mass spectrometry (MS) analysis.

### Sample Preparation for LC–MS/MS

Immunoprecipitated proteins were reduced with DTT, alkylated with iodoacetamide (both from Sigma–Aldrich), and digested overnight at 37 °C with sequencing-grade trypsin (Pierce). Peptides were then purified by solid-phase extraction using an Oasis HLB plate (Waters) and dried under vacuum. The resulting peptides were reconstituted in 0.1% formic acid and analyzed using a Q Exactive HF mass spectrometer (Thermo Fisher Scientific) coupled to an Ultimate 3000 RSLC-Nano system.

Peptides were separated on a 75 μm inner diameter, 15 cm long EasySpray analytical column (Thermo) at a flow rate of 250 nL/min. Separation was achieved using a linear gradient from 0% to 28% of buffer B over 90 minutes. Buffer A contained 2% (v/v) acetonitrile and 0.1% (v/v) formic acid in water, whereas Buffer B contained 80% (v/v) acetonitrile, 10% (v/v) trifluoroethanol, and 0.1% (v/v) formic acid in water. The mass spectrometer operated in positive ion mode with a spray voltage of 2.5 kV and ion transfer tube temperature of 275 °C. Full MS scans were acquired in the Orbitrap at a resolution of 120,000 (at m/z 200), followed by data-dependent MS/MS acquisition of up to 20 precursor ions per cycle using higher-energy collisional dissociation (HCD). Dynamic exclusion was enabled with a 20-second exclusion window for previously fragmented ions.

### Analysis of Mass Spectrometry data

Raw MS data were processed using Proteome Discoverer v3.0 (Thermo Fisher Scientific). Peptide identification was performed with Sequest HT against the UniProt Homo sapiens reference proteome. Search parameters included precursor mass tolerance of 10 ppm, fragment mass tolerance of 0.6 Da, and allowance of up to three missed cleavages. Fixed modification: carbamidomethylation of cysteine. Variable modifications: oxidation of methionine, N-terminal acetylation, loss of N-terminal methionine, methylation, dimethylation, and trimethylation of lysine and arginine, acetylation and propionylation of lysine. The false discovery rate (FDR) threshold for peptide identifications was set at 1%.

### Ileal or Colonic Crypt isolation and single cell suspension Cell Preparation

The ileum and colon was dissected from mice immediately after sacrifice. Tissues were washed five times with ice-cold phosphate-buffered saline (PBS) to remove fecal debris and luminal contents. Each sample was then opened longitudinally, cut into small fragments of approximately 2–3 cm in length, and washed three times with PBS.

To release epithelial crypts, the tissue fragments were incubated in 5 mM EDTA-PBS on ice for 30 minutes. During incubation, the tubes were gently shaken every 5 minutes to aid epithelium detachment. The resulting crypt suspensions were collected, washed once with cold PBS, and enzymatically dissociated into single cells using TrypLE Express (Invitrogen, Cat. #12604021) for 10 minutes at room temperature. Single-cell suspensions were then filtered through a 70 µm cell strainer to remove debris and undigested aggregates^10^.

### Isolation and Flow Cytometric Sorting of Intestinal Epithelial Cell Populations

Two independent experiments were performed to analyze epithelial cell populations from untreated and starved mice. In the first experiment, EpCAM⁺ /γH2AX⁺ and EpCAM⁺ /γH2AX^-^ cells were isolated from both groups.

### Antibody Labeling and Cell Sorting

For flow cytometry, cells were stained with an antibody cocktail prepared according to the experimental design. Before staining, cells were fixed with 4% paraformaldehyde (Cat. No. J19943-K2, Thermo Scientific) and subsequently permeabilized using permeabilization buffer (0.1% Triton X-100 in PBS) for 10 minutes. Fluorescently conjugated antibodies included PE–anti-EpCAM (Thermo Fisher Scientific, Cat. #12-5791-83), AF488–anti-cleaved Caspase-3 (R&D Systems/Bio-Techne, Cat. #IC835G-025), and AF647–anti-phospho-γH2AX (Ser139) (Invitrogen, Cat. #70078MP64720UG).

After staining, single-cell populations were identified and gated based on forward and side scatter to exclude doublets and debris. EpCAM⁺ epithelial cells were first gated from the total single-cell population, followed by identification of γH2AX⁺ cells within the EpCAM⁺ compartment. Subsequently, in specific experiments, cleaved caspase-3⁺ cells were gated within the EpCAM⁺γH2AX⁺ subset.

Sorting was performed using a SONY SH800S cell sorter, and purified cell populations were collected into post-sort buffer consisting of RPMI-1640 supplemented with 20% fetal bovine serum (FBS) and 15% PenStrep to maintain viability and reduce contamination. Sorted (Average 1X10^6^) cells were used immediately for downstream molecular analyses^10^.

### RNA isolation and library preparation

Total RNA form was isolated by using RNAeasy Mini Kit (Qiagen Cat# 74104) by following the manufacturer kit protocol. For RNA-seq, libraries were prepared from total RNA by using the TrueSeq RNA Library Prep Kit V2 (Illumina).

### mRNA expression by qPCR

After extracting the total RNA using RNeasy Mini Kit (Qiagen), and checking its integrity by nanodrop, the cDNA was synthesized from 1 mg of purified total RNA using High capacity cDNA Reverse Transcription Kit (Applied Biosystem). Expression of mouse and human IPMK, 18S for human was detected using suitably designed Taqman primers (Cat no. Hs00852670_g1 IPMK Invitrogen). The experiments were performed (real-time PCR Systems StepOne Plus, Applied Biosystems) in triplicate. Data were quantified for the above genes using the comparative Ct method, as described in the Assays-on-Demand User’s Manual (Applied Biosystems)^10^.

### Analysis of RNA-seq data

Sequencing reads were processed to remove adapter sequences and reads shorter than 20 bp. Reads were then aligned to either the human (Ensembl GRCh38) or mouse (Ensembl GRCm39) reference genome with STAR (v 2.7.11b; Dobin et al., 2013). To obtain read counts per gene, aligned reads were quantified using HTSeq (v 2.0.9; Putri et al., 2022) and the associated gene annotation file for the human or mouse reference genome. Differential expression analysis was performed with the R package DESeq2. Differentially expressed genes met threshold values of absolute log2 fold change greater than 1.0 and FDR < 0.05 between group comparisons^46–49^.

### Animals

All protocols were approved by the Institutional Animal Care and Use Committee (IACUC, UNLV, IACUC-01204, IACUC-01206), University of Nevada, Las Vegas. Mice were housed according to institutional guidelines, in a controlled environment at a temperature of 22°C ± 1°C, under a 12-h dark-light period, and provided with standard chow diet and water *ad libitum*. Male and female C57BL/6J and *IPMK ^F/F^*, *IPMK^IEC-/-^* (between 8-14 weeks) were used. Specifically, intestinal epithelial cells from male mice were used for biochemical and transcriptomic analysis.

### Generation of intestine-specific *IPMK* knockout mice

*IPMK^F/F^* mice were crossed with villin Cre from Jackson Laboratory (Strain #:004586) to generate intestinal epithelial cell-specific IPMK deleted mice (*IPMK^IEC-/-^*). All mice were maintained on a C57BL/6J background, and *IPMK^F/F^* were 10th-generation backcrossed. Mice were housed in a 12-hour light/12-hour dark cycle at 22°C and fed standard rodent chow. All research involving mice was approved by the University of Nevada, Las Vegas’s Institutional Animal Care and Use Committee (IACUC)^10^.

### Animal Fasting Experiment

C57BL/6J wild-type, *IPMK^F/F^*, and *IPMK^IEC-/-^* mice will undergo a 12-hour overnight fasting period with free access to water under standard housing conditions. After fasting, mice will be euthanized, and tissues will be collected for histopathological and biochemical analyses. For the refeeding group, mice will be provided with food for four hours following the 12-hour overnight fasting. Subsequently, the mice will be euthanized, and tissues will be harvested for histopathological and biochemical analyses.

### Intestinal cell isolation

The ileum was dissected from mice. After five times washing with ice-cold PBS, the tissues were opened longitudinally and cut into small fragments (2–3 cm in length). To prepare single-cell suspension after washing three times with PBS, small intestinal fragments were incubated in 5 mM EDTA-PBS on ice for 30 min, during which the tissues were shaken vigorously every 5 min. Isolated epithelial cells and crypts were washed five times with cold PBS. Cell suspensions were passed through a 70-mm cell strainer after that used for experiment^10^.

### EdU Staining

To assess DNA synthesis as an indicator of epithelial regeneration, mice received a single intraperitoneal injection of 5-ethynyl-2′-deoxyuridine (EdU) at a dose of 200 µg/kg in PBS. Two hours post-injection, the mice were euthanized, and small intestines were harvested and processed for histological analysis. Detection of EdU incorporation was performed using the Click-iT™ EdU Cell Proliferation Kit for Imaging, Alexa Fluor™ 488 (Thermo Fisher), following the manufacturer’s instructions^50^.

### Histology and Immunohistochemistry

Mice were humanely euthanized by controlled exposure to CO₂, following institutional ethical guidelines for animal care and use. Immediately after euthanasia, the intestine and colon were carefully excised and thoroughly flushed with cold phosphate-buffered saline (PBS) to remove any residual luminal contents. Following this, the tissues were gently stretched to their full length and wrapped longitudinally in filter paper to maintain their orientation and structural integrity during fixation. The wrapped tissues were then immersed in 10% phosphate-buffered paraformaldehyde (PFA) and fixed for an appropriate period to preserve tissue morphology and architecture. After fixation, the intestinal tissues were Swiss-rolled, a method that allows the entire length of the intestine to be visualized in a single tissue section. The Swiss-rolled tissues were subsequently processed through 70%, 90%, and 100% ethanol, followed by treatment with xylene. Dehydrated tissues were then embedded in low-melting-point paraffin wax (65°C) to prepare tissue blocks. Six-micron-thick sections were cut using a microtome and mounted onto Mayer’s albumin-coated glass slides to ensure proper adhesion of sections.

For immunofluorescence staining, paraffin-embedded sections were deparaffinized using Histo-Clear (Thermo Fisher) and rehydrated through successive washes of 100%, 90%, and 75% ethanol, followed by DI water. Heat-induced epitope retrieval (HIER) was performed using citrate buffer for one hour at 80°C, ensuring unmasking of antigenic sites. To minimize non-specific binding, sections were blocked with filtered 3% goat serum for 1 hour at 37°C.

Primary antibodies, prepared at concentrations recommended by the manufacturer, were diluted in a blocking solution containing 1% Triton X-100, and applied to tissue sections. Slides were incubated overnight at 4°C in a humidified chamber to allow for specific antigen-antibody interactions. The following day, sections were washed thoroughly with TBST (Tris-buffered saline with Tween-20) and incubated with appropriate fluorescently labeled secondary antibodies for 60 minutes at 37°C. After five consecutive washes with TBST to remove unbound antibodies, sections were counterstained with DAPI (1 μg/mL in PBS) to visualize nuclei.

Finally, stained slides were imaged using a Zeiss Confocal microscope (LSM 800), and digital images were captured for analysis. Signal intensities were quantified using ImageJ software^50^.

### Mouse Intestinal Organoid Culture

Proximal small intestines (∼15 cm) were dissected from mice immediately after sacrifice under sterile conditions. The lumen was flushed with ice-cold PBS using a 10-ml syringe fitted with a 21G needle to remove luminal contents. Intestines were opened longitudinally, cut into ∼5-mm pieces, and transferred to a 50-ml conical tube containing 25 ml ice-cold PBS. Tissues were washed by inversion and PBS replacement repeatedly until the supernatant appeared clear.

For crypt isolation, tissue fragments were incubated in 10 ml ice-cold 5 mM EDTA in PBS and vigorously triturated with a 10-ml serological pipette. After allowing fragments to settle, the supernatant was removed and replaced with fresh 5 mM EDTA-PBS and incubated on a benchtop roller at 4 °C for 10 min. This step was repeated with 5 mM EDTA-PBS for an additional 30 min at 4 °C. Fragments were then washed with cold PBS. Crypts were released by adding PBS containing 0.1% BSA and shaking vigorously; successive supernatant fractions were collected, and the crypt-enriched fractions were identified microscopically.

The crypt-rich fraction was mixed 1:1 with basal medium supplemented with DNase I (15 U/ml), passed through a 70-µm cell strainer into a 1% BSA-coated 50-ml tube, and centrifuged at 300 × g for 5 min at 4 °C. The pellet was resuspended in basal medium containing 5% FBS and centrifuged at 100 × g for 5 min to enrich intact crypts. Crypts were finally resuspended in a small volume of basal medium, counted, and adjusted to yield approximately 100-500 crypts per Matrigel dome.

For organoid culture, isolated crypts were resuspended in IntestiCult™ Organoid Growth Medium and mixed 1:1 with undiluted, ice-cold Matrigel. Aliquots of 50 µl of the crypt–Matrigel suspension were plated as domes into pre-warmed 48-well plates and incubated at 37 °C for 10 min to allow Matrigel polymerization. Following solidification, 500 µl of pre-warmed IntestiCult medium was gently added to each well. Organoids were maintained at 37 °C in a humidified incubator with 5% CO₂, and medium was replaced every 2–3 days until organoids reached the desired stage for downstream experiments^50^.

### Statistical analysis

All plots and statistical analyses were performed with Prism 9 (GraphPad) software. Statistical significance was determined by either Student’s t-test (two-tailed) for two groups or 1-way ANOVA for multiple groups with similar samples. Error bars represent the standard deviation of the mean, and dots indicate replicates or the number of animals employed. Results were representative of at least three independent experiments (n). Graphs were generated by using either Prism 9 (GraphPad) software or R software.

## Reporting summary

Further information on research design is available in the Reporting Summary linked to this article.

## Data availability

RNA-seq and CUT&RUN sequencing datasets generated in this study have been deposited in the SRA database under accession code **PRJNA1451917**.

## Acknowledgements

This work was supported by 5P20GM121325 COBRE grants and University of Nevada, Las Vegas start-up funds to Prasun Guha. We like to thank NIPM’s Genomic Core for assisting with instruments and experiments. We would like to thank Seungman Park from the University of Nevada, Las Vegas for providing the MCF7 cell line; Hafiz Ahmed from the University of Maryland for providing the B16F10, MDAMB231, and HepG2 cell lines; and Xiaoyu Zhang, Northwestern University, USA, for providing the *FBXO22* KO A549 cell line. We would like to thank Daolin Tang from UT Southwestern Medical Center for generously providing MEF *WT and ATG5* KO cells. We would like to thank Mark Donowitz, Douglas Robinson, Ted Dawson, and Jeremy Nathans from Johns Hopkins Medicine and Matthew Ciorba from Washington University for their careful review and valuable suggestions that improved this manuscript.

## Competing interests

The authors declare no conflict of interest. All authors reviewed the results and approved the final version of the manuscript.

## Author Contributions

PG conceived the study. PG and SC designed the experiments. SC performed most of the experiments. LVP, ZS, PG, TT, and NT independently performed and validated major biochemical experiments. SC performed wet-lab experiments related to NGS studies. RV, ZS, MH, and LVP performed most of the NGS data analysis. FB, KR, and HJJ generated the cell-permeable InsP_6_. NT and SC performed animal model and microscopy-related experiments and image acquisition. LG and MW helped in doing confocal microscopy. TT, RM, KS, and GK analyzed microscopy-related raw data and performed analysis. AS and HJJ analyzed mass spectrometry-based inositol profiling from cells and tissue. FB and FJD developed and characterized HDAC3 PROTAC. SC, RV, and ZS designed the figures. KH, RR, AG, MG, CHC, and PD provided and analyzed human patient samples. PG and SC wrote the manuscript. SC takes responsibility for all wet lab data.

## Declaration of generative AI and AI-assisted technologies in the manuscript preparation process

During the preparation of this manuscript, the authors used ChatGPT (v5.2) and Perplexity to assist with language refinement and readability. The authors reviewed and edited all generated content and accept full responsibility for the final manuscript.

---

**Supplementary Figure 1. Short term fasting selectively trigger caspase 3 activation in pre-existing DNA-damged cells.**

**(A)** Immunoblot and densitometric analysis of p-γ-H2AX and cleaved caspase-3 in untreated (UT) and starved conditions (n = 3; mean ± SD). (**B)** After flow purification, quantification of γ-H2AX⁺ cell populations in untreated and starved groups (n = 3; mean ± SD). **(C)** Immunoblot followed by densitometric analysis of p-γ-H2AX (Ser139) in the ileal crypt showed that 12-hour starvation reduced p-γ-H2AX (Ser139) levels (n = 15). **(D)** Hematoxylin and eosin staining of colonic sections from untreated and starved mice; scale bar, 100 μm. **(E)** Time-course immunoblot at indicated starvation time points (0–12 h) with densitometric analysis (n = 3 for 0-10hr,n=7 for 12hr; mean ± SD). **(F)** Schematic of short-term food restriction (6 h) prior to analysis. Created in BioRender (CHATTERJEE, S., 2026, CHATTERJEE, S. (2026) https://BioRender.com/mtrovit.). **(G)** Immunoblot and quantification from EpCAM⁺/p-γ-H2AX⁺ and EpCAM⁺/p-γ-H2AX⁻ cells under untreated and 6 hr starved conditions (n = 3; mean ± SD). **(H)** Hematoxylin and eosin staining of DSS-treated and starved+DSS colon tissue; scale bar, 100 μm.

**Supplementary Figure 2. CDK5RAP3 induction during fasting suppresses RPL26 UFMylation**

**(A)** Immunoblot analysis of RPL26 UFMylation (UFM1 and UFM2 conjugates) in cells expressing GFP- or myc-tagged CDK5RAP3, with corresponding CDK5RAP3, CHOP, IRE1, calreticulin, and actin controls. Immunoblot (IB) with either RPL26 or UFM1 antibody to validate the UFMylation.(n=3)

**(B)** Time-course immunoblot of CDK5RAP3, CHOP, IRE1, and RPL26 UFMylation during starvation (0–12 h) (n = 3).

**(C)** Immunoblot analysis comparing EpCAM⁺/p-γ-H2AX⁺ and EpCAM⁺/p-γ-H2AX⁻ cells under indicated 6hr starved conditions. (n = 3).

**(D)** Immunoblot showing p-γ-H2AX, total H2AX, cleaved caspase-3, and GFP in cells expressing GFP-CDK5RAP3 under untreated (UT) and 60 min UV treated conditions. (n = 3).

**(E)** Immunoblot analysis of CDK5RAP3, CHOP, and IRE1 in liver, lung, and pancreas from untreated and starved mice (n = 3).

**(F)** Genome browser tracks showing H3K27ac distribution across the CDK5RAP3 locus on chromosome 11 in untreated (blue) and starved (orange) conditions. Yellow highlights show enrichment in starved.

**Supplementary Figure 3. Fasting does not alter p300 HAT activity.**

(A) P300 immunopurified from untreated (UT) and starved mice IEC followed by activity assay in vitro. Western blot showing total protein level. (n = 3;mean ± SD).

(B) Anti-CDK5RAP3 immunoprecipitation from 100μg, 400μg, 800μg followed by CDKRAP3 immunoblotting showing equal amount of CDK5RAP3 immunopurified from 400μg of protein. (n=3)

**Supplementary Figure 4. Growth factor deprivation modulates histone acetylation, HDAC3 activity, and CDK5RAP3-associated ER stress.**

**(A)** Schematic of the SILAC-based proteomic workflow used to quantify histone acetylation. Created in BioRender. CHATTERJEE, S. (2026) https://BioRender.com/mtrovit.

**(B)** Mass spectrometry analysis of histone acetylation sites under GFD from HCT116 (n = 3;mean ± SD).

**(C)** Immunoblot analysis of H3K9Ac, H3K27Ac, and H4K16Ac in GFD-treated HCT116 cells; SAHA is pan HDAC inhibitor (n = 3).

**(D)** HDAC3 immunoprecipitation from 400 μg of protein lysate and western blot of CDK5RAP3 and other proteins from HCT116 (n = 3).

**(E)** CDK5RAP3 immunoprecipitation from 400 μg of protein lysate and western blot of P300 from HCT116 (n = 3).

**(F)** HDAC activity assays for HDAC1/2, HDAC3, and HDAC8 from HCT116 cells under untreated (UT) and GFD (n = 3; mean ± SD).

**(G)** Immunoprecipitation of HDAC1, HDAC2, HDAC3, and HDAC8 from equal lysate input with corresponding lysate and IP fractions (n = 3).

**(H)** HDAC3 activity assays from B16F10, A549, MDA-MB231, HepG2, MCF7, MEF, HEK, and Caco2 cells under untreated (UT) and GFD (n = 3; mean ± SD).

**(I)** Immunoblot analysis of HDAC3 protein levels in indicated cell lines under indicated conditions (n = 3).

**(J)** Immunoblot analysis of CDK5RAP3 (yellow Proteintech, green Bethyl), RPL26 UFMylation, CHOP, and IRE1 in HDAC3 knockout HCT116 cells in indicated conditions (n = 3). Used Proteintech and Bethyl antibodies to validate CDK5RAP3 expression.

**(K)** Transient reconstitution of *HDAC3* knockout cells with wild-type or InsP₆-binding–defective HDAC3 (HDAC3^HOIP^) plasmid (n = 3) followed by western blot.(n = 3)

**(L)** Co-immunoprecipitation of HDAC3 with SMRT and NCoR under indicated conditions (n = 3).

**(M)** Co-immunoprecipitation of HDAC3 with FLAG-tagged SMRT deacetylase-activating domain (FLAG-DAD) with or without 50μM CP-InsP₆ treatment for 24hr (n = 3).

**(N)** Quantification of intracellular inositol phosphates by mass spectrometry under GFD (n = 3). Untreated (UT), GFD.

**(O)** RT–qPCR analysis of IPMK mRNA expression under Untreated (UT) and GFD (n = 3; mean ± SD).

**(P)** Immunoblot analysis of CDK5RAP3, CHOP, IRE1, and RPL26 UFMylation following CP-InsP₆ treatment (n = 3).

**Supplementary Figure 5. FBXO2 depletion does not rescue IPMK degradation during growth factor deprivation**

**(A)** Immunoblot analysis of IPMK following FBXO2 knockdown in untreated and GFD HCT116 cells. (n=3).

**Supplementary Figure 6. Acetylation of CDK5RAP3 prevents its autophagic degradation.**

**(A)** Immunoblot analysis of CDK5RAP3 in HCT116 cells treated with the autophagy inhibitor bafilomycin A1 or the proteasome inhibitor bortezomib under GFD or untreated conditions. (n = 3).

**(B)** Immunoblot analysis of CDK5RAP3 WT and *ATG5 KO* MEF cells under basal conditions. (n = 5).

**(C)** Immunoblot analysis of CDK5RAP3 following treatment with the ULK1/2 inhibitor SBI-0206965 (10 μM for 24 hr) in HCT116 cells under basal conditions. (n = 3).

**(D)** Immunoblot analysis of CDK5RAP3 in cells under GFD or untreated conditions with CP-InsP₆, the p300 inhibitor A485, or the autophagy inhibitor bafilomycin A1. (n = 3).

**Supplementary Figure 7. TGFβ specifically restores the IPMK–HDAC3 axis during growth factor deprivation**

**(A)** HDAC3 enzymatic activity measured in MEF cells under untreated (UT), GFD, and growth factor–stimulated conditions (TGFβ, IGF, VEGF, EGF, FGF, PDGF). (n = 3; mean ± SD).

**(B)** Immunoblot analysis of H3K9ac, H3K27ac, H4K16ac, and HDAC3 in untreated, GFD, and TGFβ-stimulated MEF cells. (n = 3).

**(C)** Immunoblot analysis of CDK5RAP3, RPL26 UFMylation, CHOP, IRE1, and phosphorylated SMAD2 in serum-fed HCT116 cells treated with the TGFβ receptor inhibitor RepSox. (n = 3).

**(D)** RepSox treatment increased CDK5RAP3 acetylation immunoprecipitation from 400 μg protein. (n=3)

**(E)** Quantification of HDAC3 enzymatic activity in MEF cells under GFD with TGFβ stimulation or growth factor cocktail excluding TGFβ (PDGF+EGF+VEGF+IGF+FGF). (n = 3; mean ± SD).

**(F)** Immunoblot analysis of IPMK, CDK5RAP3, CHOP, and IRE1 in MEF cells under GFD with TGFβ stimulation, RepSox treatment, or growth factor cocktail (PDGF+EGF+VEGF+IGF+FGF)-TGFβ (n = 3).

**Supplementary Figure 8. CP-InsP_6_ effects in enteroids.**

**(A)** Representative bright-field images of intestinal enteroids. Scale bar 60µm.

**(B)** Enteroids co-treated with CP-InsP_6_ and Rexpsox or CP-InsP6 and HDAC3-PROTAC followed by western blot.(n=3).

## Notes

### Competing Interest Statement

The authors have declared no competing interest.

