## Supplemental figure for "Fasting disrupts the InsP₆–HDAC3 axis to drive ER stress–mediated clearance of DNA-damaged cells and enforce tissue quality control"

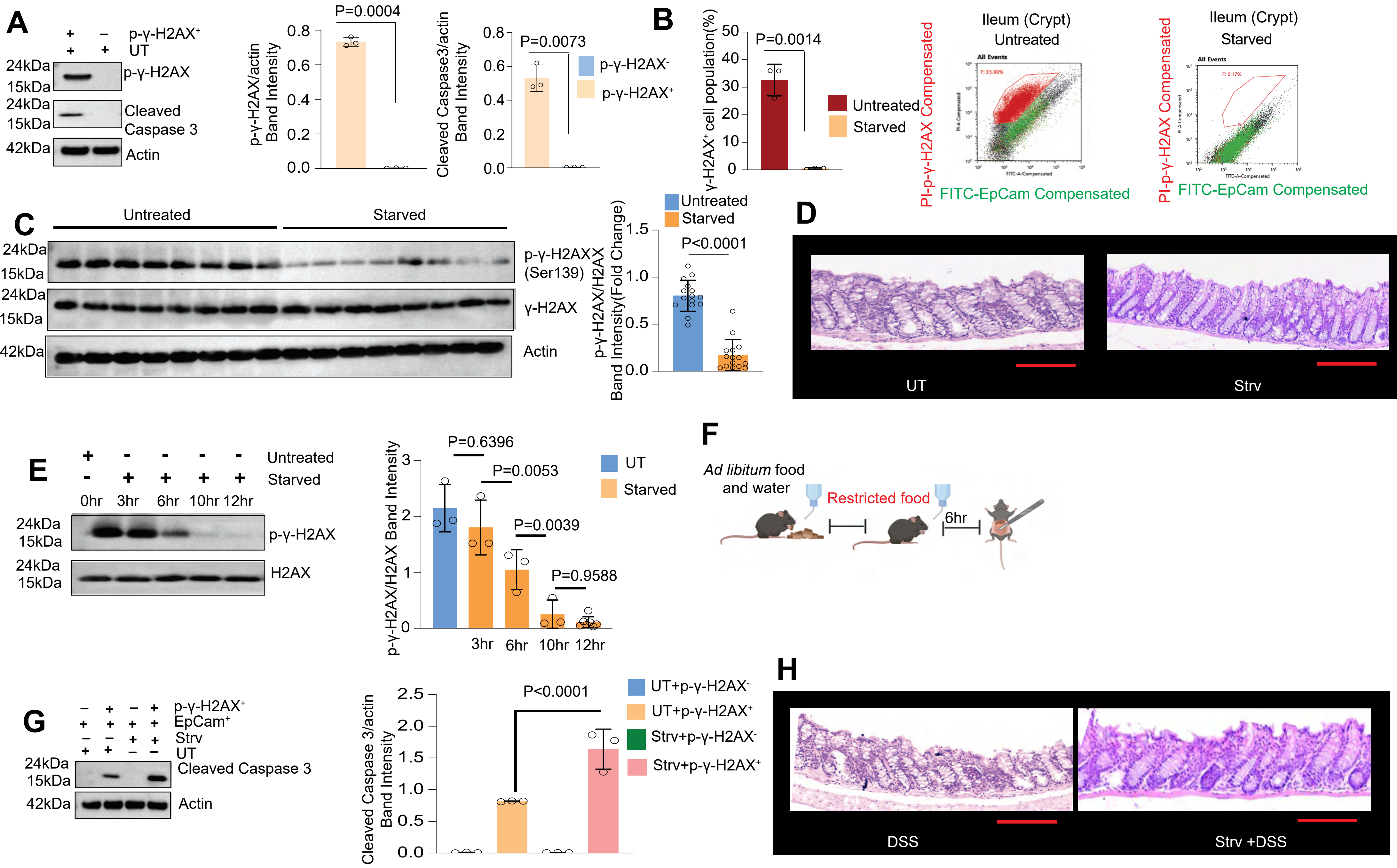


**Supplementary Figure 1**

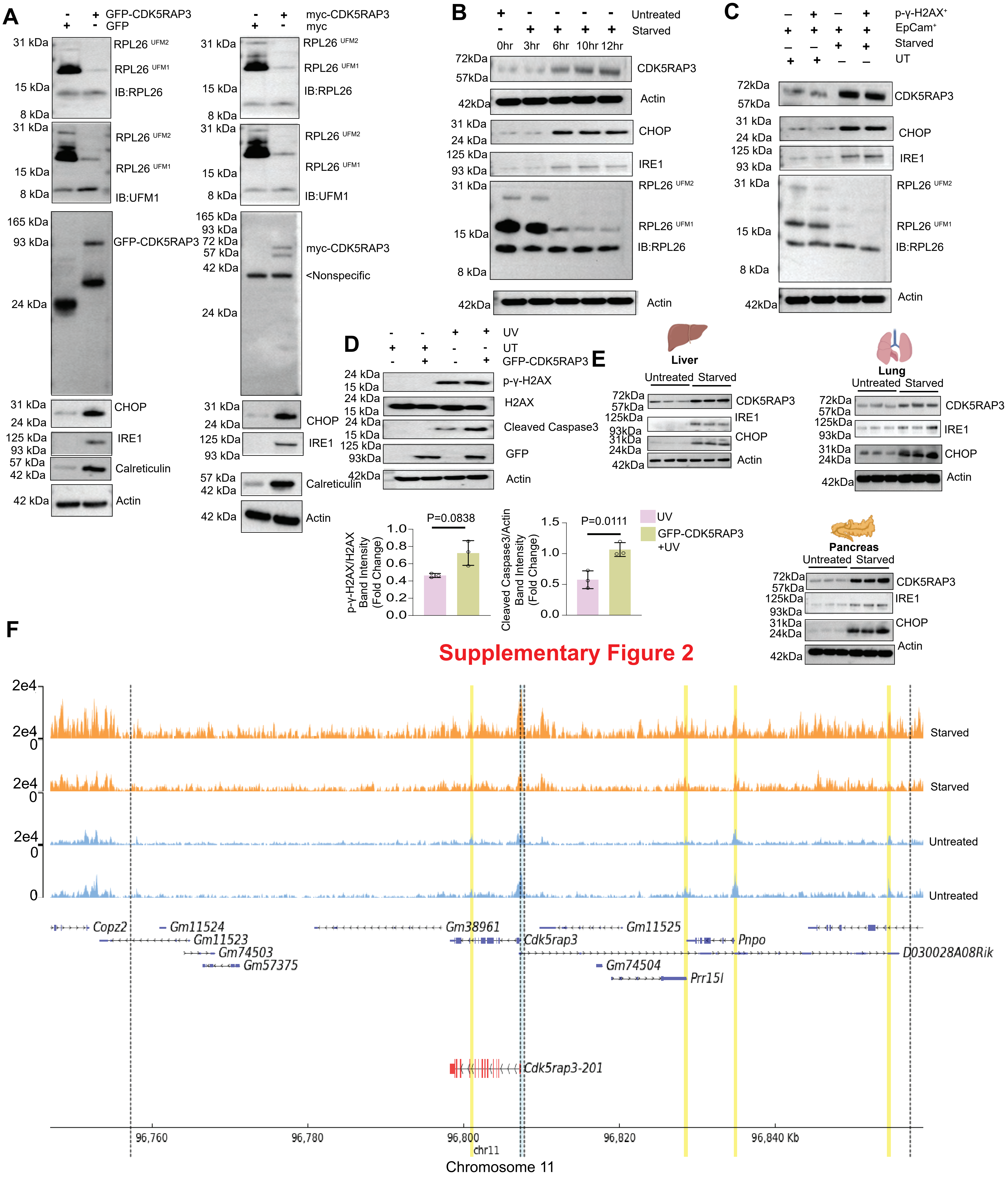


#### Supplementary Figure 2

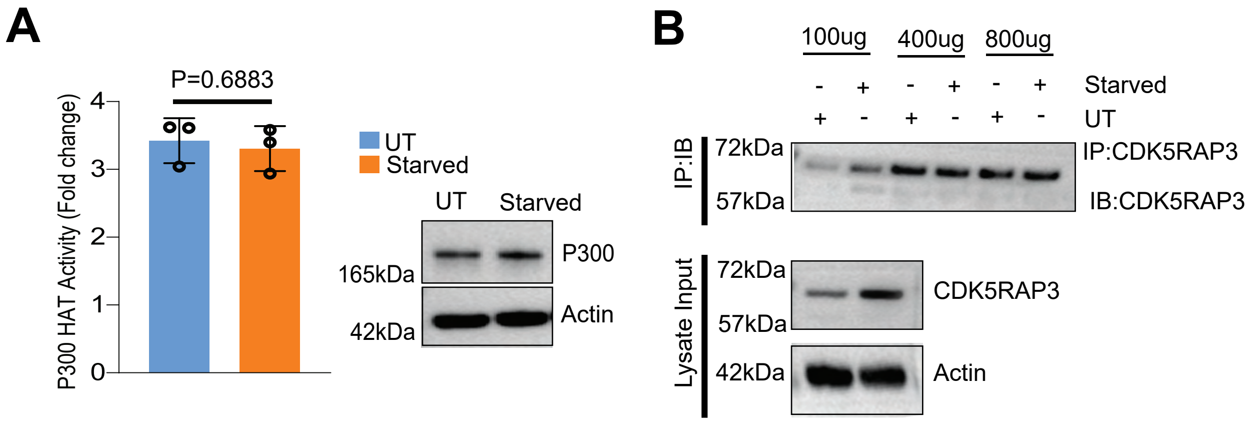


### Supplementary Figure 3

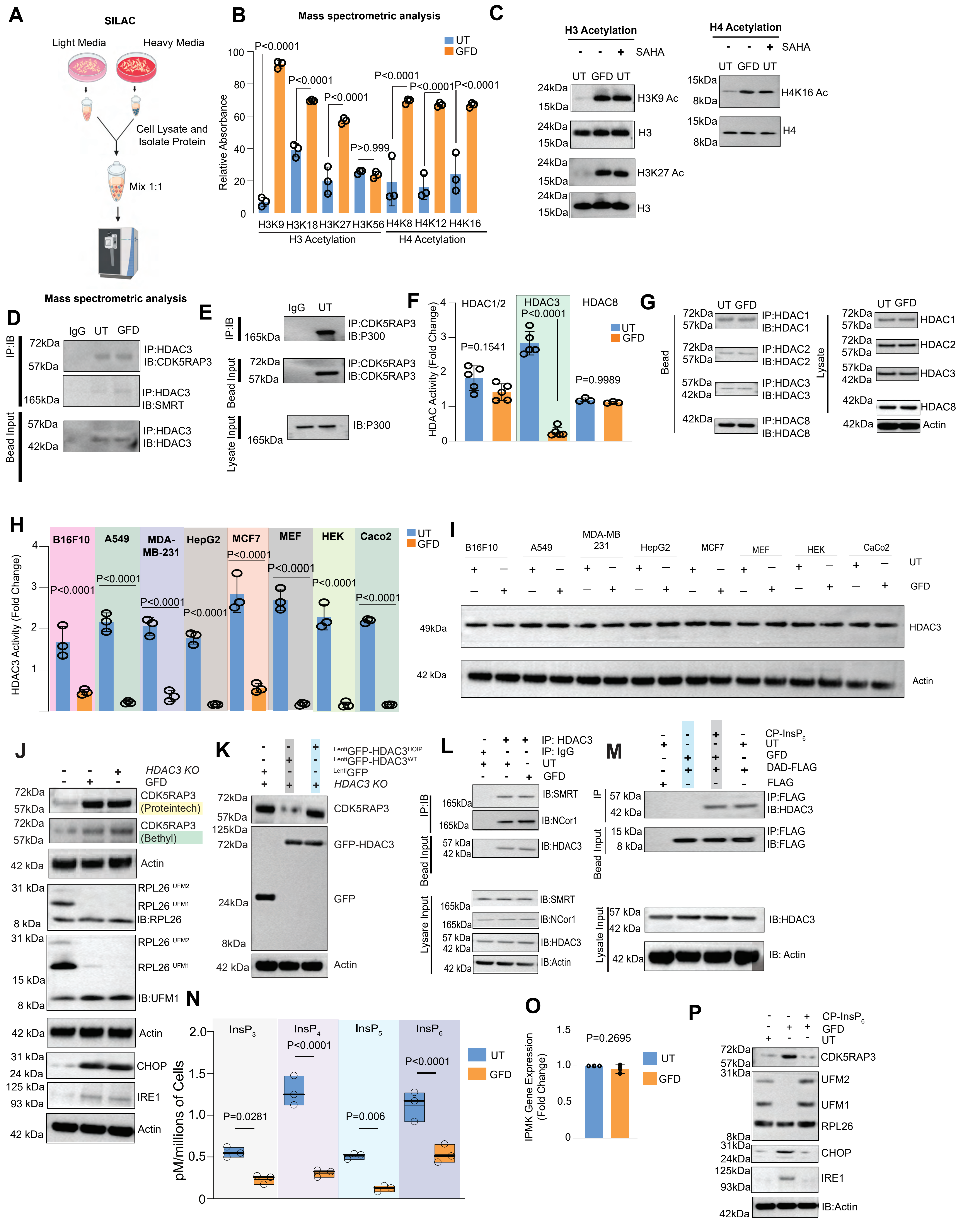


**Supplementary Figure 4**

A

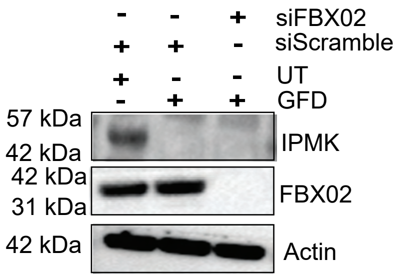


**Supplementary Figure 5**

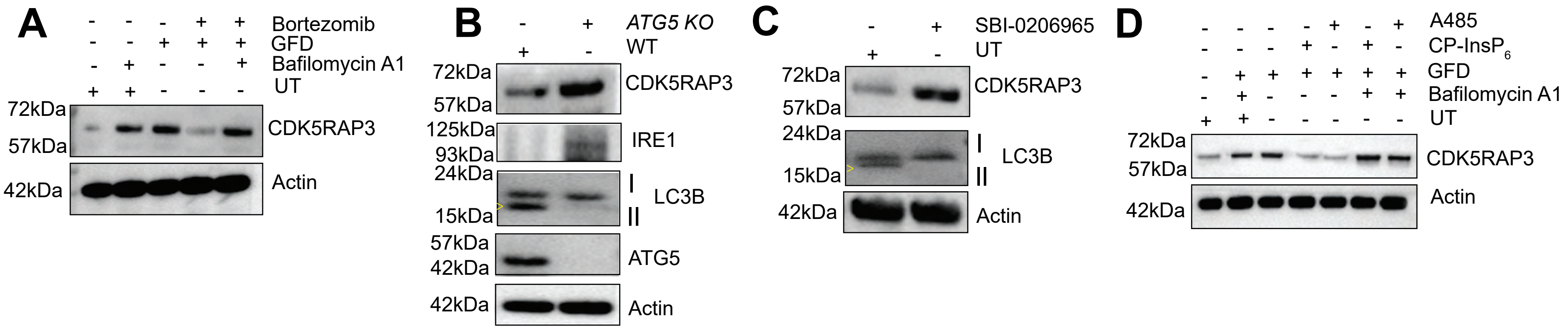


**Supplementary Figure 6**

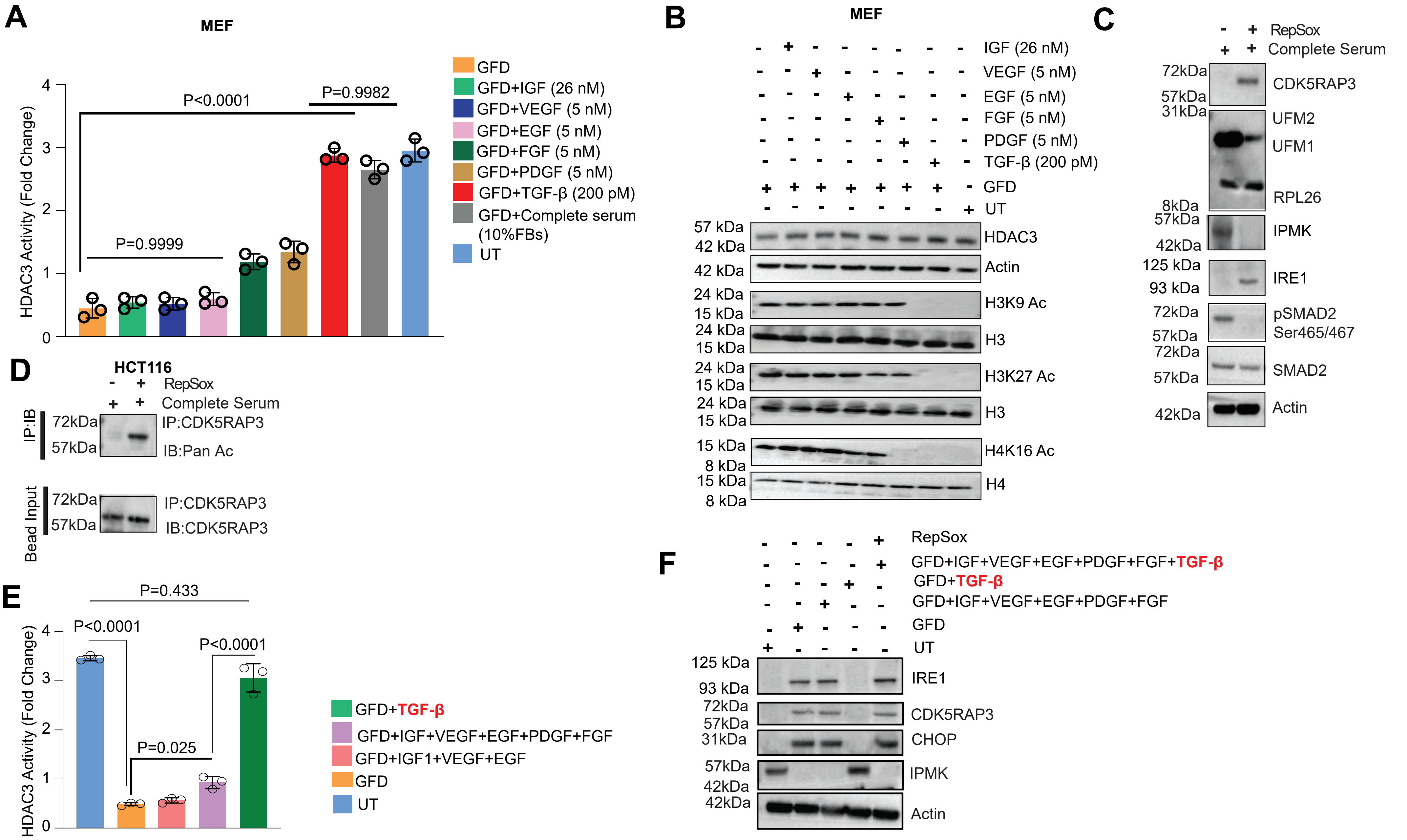


Supplementary Figure 7

A

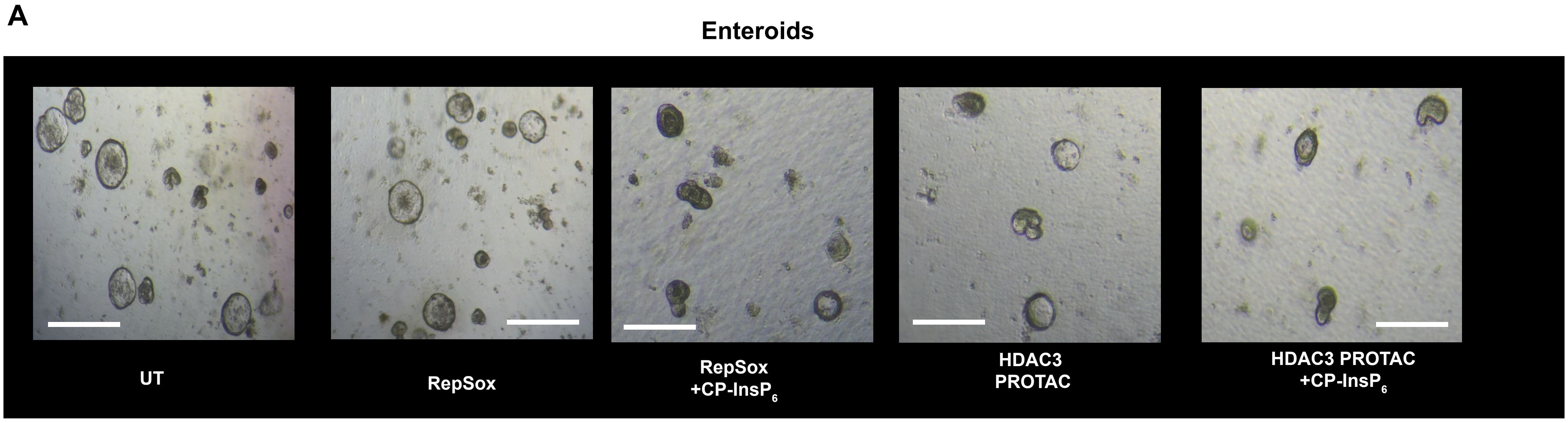


B

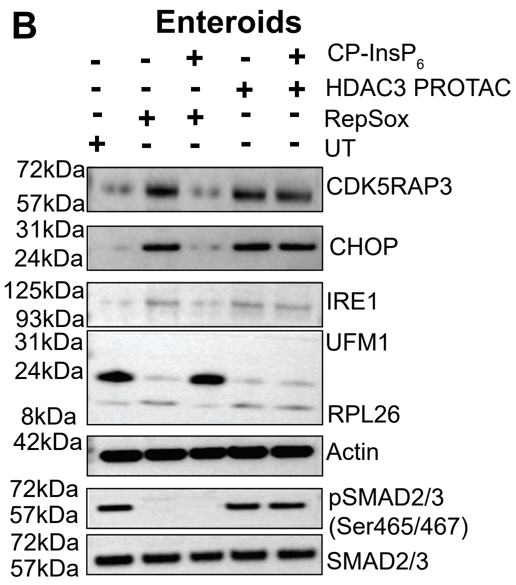


Supplementary Figure 8
